# MASCOT-Skyline integrates population and migration dynamics to enhance phylogeographic reconstructions

**DOI:** 10.1101/2024.03.06.583734

**Authors:** Nicola F. Müller, Remco R. Bouckaert, Chieh-Hsi Wu, Trevor Bedford

## Abstract

The spread of infectious diseases is shaped by spatial and temporal aspects, such as host population structure or changes in the transmission rate or number of infected individuals over time. These spatiotemporal dynamics are imprinted in the genome of pathogens and can be recovered from those genomes using phylodynamics methods. However, phylodynamic methods typically quantify either the temporal or spatial transmission dynamics, which leads to unclear biases, as one can potentially not be inferred without the other. Here, we address this challenge by introducing a structured coalescent skyline approach, MASCOT-Skyline that allows us to jointly infer spatial and temporal transmission dynamics of infectious diseases using Markov chain Monte Carlo inference. To do so, we model the effective population size dynamics in different locations using a non-parametric function, allowing us to approximate a range of population size dynamics. We show, using a range of different viral outbreak datasets, potential issues with phylogeographic methods. We then use these viral datasets to motivate simulations of outbreaks that illuminate the nature of biases present in the different phylogeographic methods. We show that spatial and temporal dynamics should be modeled jointly even if one seeks to recover just one of the two. Further, we showcase conditions under which we can expect phylogeographic analyses to be biased, particularly different subsampling approaches, as well as provide recommendations of when we can expect them to perform well. We implemented MASCOT-Skyline as part of the open-source software package MASCOT for the Bayesian phylodynamics platform BEAST2.

## Introduction

Infectious diseases are a major burden on public health systems around the world (Vos *et al*., 2020). Different data sources and methods exist to understand how these diseases spread quantitatively. Mainly, this relies on case data, that is, counts of when and where cases of a particular disease occurred. However, given case counts suffer from various limitations, including under-ascertainment, delays in reporting, and changes in the rate of under-ascertainment over time and between locations (Gibbons *et al*., 2014), there is continued interest in alternative data sources.

One such data source, genomic data, is increasingly being collected for infectious disease surveillance (Gardy and Loman, 2018), though substantial differences in genomic surveillance exist across the globe (Brito *et al*., 2022). Genomic data can be obtained by sequencing a subset of laboratory-confirmed cases. Pathogen genomes can give us a window into how diseases spread. While pathogens are transmitted between individuals, random mutations to their genomes accrue over time. These random changes to their genomes can then be used to reconstruct the relatedness of viruses sequenced from individuals. The evolutionary relationship, or the phylogenetic tree, of the pathogens approximates the transmission history linking these individuals. From this phylogenetic tree, one can infer the transmission dynamics of infectious diseases using phylodynamic methods even if only a subset of individuals in the transmission history is sequenced (Grenfell *et al*., 2004). Phylodynamic methods utilize the branching patterns of timed phylogenetic trees to learn about the underlying population dynamics that created them (Holmes and Grenfell, 2009; Volz *et al*., 2013). This information can be inferred using forwards-in-time birth-death (Kendall, 1948) or backwards-in-time coalescent models (Kingman, 1982). Birth-death models describe how lineages multiply (birth), go extinct (death), and are sampled. The birth and death rates and their changes over time can be used to describe the transmission rates, becoming uninfectious rates, or effective reproduction numbers (Stadler *et al*., 2013). Coalescent models, however, describe how lineages coalesce in the past, meaning when they share a common ancestor. The rates at which two random lineages and a population share a common ancestor are lower if the population is larger and vice versa. The coalescent is typically parameterized by the effective population size (*Ne*), which is proportional to the number of infected individuals in a population and inversely proportional to the transmission rate in that population (Volz *et al*., 2009; Volz, 2012). In contrast to case-based inference methods and birth-death methods, coalescent approaches infer population size dynamics from the relatedness of cases instead of the dynamics in the number of samples. Nonetheless, they can still suffer somewhat from biases under specific sampling assumption (Karcher *et al*., 2016).

By modeling changes in the effective population size over time Ne(t), coalescent approaches can be used to model changes in pathogen prevalence or generation time over time. One can use deterministic parametric approaches to model changes in the population sizes over time (Volz *et al*., 2009) or simulate population trajectories from stochastic compartmental models (Popinga *et al*., 2015). Alternatively, non-parametric approaches, typically called skyline models, can be used (Strimmer and Pybus, 2001). These methods allow the effective population sizes to vary over time in a piecewise, constant fashion. Different skyline approaches vary in how changes in effective population sizes are parameterized. Some a priori assume the number of change points to be fixed allows the effective population size to change at coalescent events (Drummond *et al*., 2005; Minin *et al*., 2008; Bouckaert, 2022). Others, typically called skygrid methods, allow the effective population sizes to vary at pre-determined points in time (Gill *et al*., 2013) or split the height of the tree into equally sized epochs (Bouckaert, 2022). Coalescent models have been previously deployed to, for example, study the change in the prevalence of hepatitis C (Pybus *et al*., 2003), seasonal influenza (Rambaut *et al*., 2008) and tuberculosis (Merker *et al*., 2015).

A further advantage of inferring transmission dynamics from genomic data is that we can learn about how cases between locations are connected. We can use this information to infer spatial transmission dynamics, which are not readily accessible from occurrence data alone. A small set of examples for this work includes studies on the early spread of HIV (Faria *et al*., 2014; Worobey *et al*., 2016), the global circulation of seasonal influenza (Bedford *et al*., 2015), and the cross-species transmission of MERS coronaviruses (Dudas *et al*., 2018) or yellow fever (Faria *et al*., 2018). Related approaches can be used in “who infected who” approaches that seek to determine transmission directionality between individuals (see, for example, De Maio *et al*. (2016)), showing the broad range of applications of methods that model population structure.

Different methods exist to do so, including discrete trait analyses (DTA) (Lemey *et al*., 2009), structured birth-death (Maddison *et al*., 2007; Stadler and Bonhoeffer, 2013; Kühnert *et al*., 2016), or structured coalescent methods (Takahata, 1988; Hudson *et al*., 1990; Notohara, 1990). DTA is conceptually different from structured birth-death and coalescent models. DTA only models the movement of viral lineages without explicitly modeling anything about branching processes. They are, therefore, also referred to as neutral trait models, meaning that they model the evolution of a trait, such as geographic location, on top of an existing phylogenetic tree. DTA has arguably been the most popular method of the here described methods, partly due to its ease of use and computational speed. However, biased sampling in DTA models can often lead to biased model results (De Maio *et al*., 2015). Structured birth-death models describe the birth, death, sampling, and movement of lineages between discrete sub-populations or demes forward in time.

Structured coalescent models model how lineages share a common ancestor within and move between sub-populations, from present to past, backward in time. The structured coalescent is parameterized by effective population size (Ne) and migration rates, which can be related to epidemiologically more meaningful parameters, such as the prevalence and transmission rates (Volz, 2012). Structured coalescent methods largely assume that the rates of coalescence and migration are constant over time, though deterministic approaches to model parametric dynamics from compartmental models exist (Volz and Siveroni, 2018) While structured coalescent approaches are historically not used as frequently as discrete trait analyses, there are some distinct advantages to these types of methods, including potentially being less subject to sampling biases (De Maio *et al*., 2015), while still being able to analyze larger datasets (Müller *et al*., 2018). One of the limiting factors of structured coalescent methods is their assumption of populations to be constant over time. This assumption is, however, rarely appropriate and can lead to the biased reconstruction of the within-deme and the between-deme dynamics (Layan *et al*., 2023).

Here, we introduce a phylodynamic framework to infer non-parametric effective population size (Ne) dynamics under the marginal approximation of the structured coalescent MASCOT (Müller *et al*., 2018). The effective population sizes are estimated at predefined points in time, between which we assume exponential growth dynamics (Volz and Didelot, 2018). As such, we allow the Ne’s to continuously change over time instead of assuming piecewise constant dynamics, as is typically used in skyline approaches (for example Gill *et al*. (2013)). We use a Gaussian Markov Random Field (GMRF), as in Gill *et al*. (2013) for unstructured populations, to model the temporal correlation between Ne’s. We then estimate Ne trajectories for each sub-population in the model using Markov chain Monte Carlo (MCMC) by using MCMC operations that learn the correlation structure between the different parameters Baele *et al*. (2017). We first show, using simulations, that we can retrieve non-parametric population dynamics and migration rates of different sub-populations from phylogenetic trees. We then show how accounting for population structure improves the inference of population dynamics and vice versa. Lastly, we compare the ancestral state reconstruction and inference results of migration rates between MASCOT-Skyline and DTA (Lemey *et al*., 2009) using a dataset of SARS-CoV-2 sequences and Susceptible-Infected-Recovered (SIR) simulations. We implemented MASCOT-Skyline as part of the BEAST2 package MASCOT (Müller *et al*., 2018), a package for the Bayesian phylogenetics software platform BEAST2 (Bouckaert *et al*., 2019).

## Results

### Nonparametric population dynamics and migration patterns can be recovered from phylogenetic trees

We first performed a well-calibrated simulation study using a two-state structured coalescent model in MASTER (Vaughan and Drummond, 2013), to validate the ability of MASCOT-Skyline to retrieve non-parametric population size dynamics. We simulated effective population size trajectories from a Gaussian Markov random field (GMRF). We sampled the natural logarithm of the effective population size at time *t* = 0 in state *a ln*(*Ne_a_*(*t* = 0)) from a normal distribution 𝒩 (0, 1). For each Ne at time *n* > 0, we sampled the *Ne* from *ln*(*Ne*(*t* = *n*)) ~ 𝒩 (*ln*(*Ne*(*t* = *n* − 1)), 0.5). Between adjacent Ne’s, we assume exponential growth. We repeated this to get the Ne trajectories of both states. We then sample the forward-in-time migration rates between the two states from an exponential distribution with a mean of 1. We compute the backward-in-time migration rates over time from the forward migration rates and the *Ne* trajectories using equation 1. Next, we simulate one phylogenetic tree using 800 leaves, 400 from each location, and infer the *Ne* trajectories and migration rates using MASCOT-Skyline from that tree. We use an exponential distribution with a mean of 1 for the migration prior and the above specification of the GMRF for the Ne prior. We repeated this process 100 times.

In Figure 1, we show, for four of the total 100 randomly chosen replicates, that MASCOT-skyline can retrieve these nonparametric population dynamics from phylogenetic trees. Using these simulations, we obtain a 94% coverage of the 95% highest posterior density interval (HPD) of the true Ne value (see Figure S1A). The forward-in-time migration rates are also recovered well by MASCOT-Skyline (see Figure S1B), though, at 89%, the coverage is below the expected range (91% to 99%) of coverage estimates for 100 replicates. This is not unexpected as MASCOT is an approximation of the structured coalescent (Müller *et al*., 2017).

**Figure 1:**
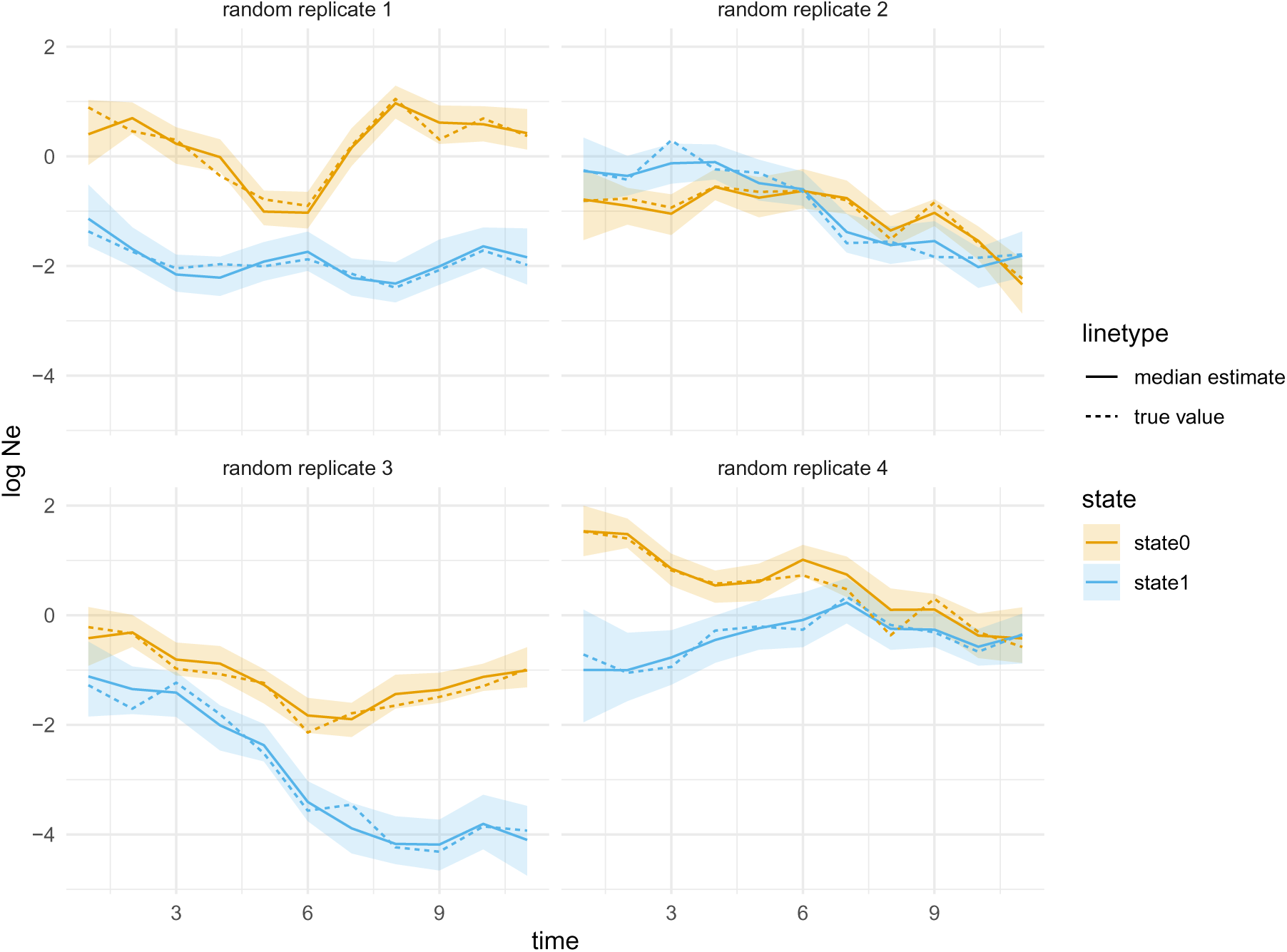
Inferred effective population size trajectories from simulated data. Here, we show the inferred effective population size dynamics with the line denoting the median inferred log Ne’s. The shaded areas denote the bounds of the 95% highest posterior density interval. The plots show the results for four of the 100 replicates chosen randomly.

### Assumptions about the population dynamics drive ancestral state reconstruction in structured coalescent models

Spatial and temporal population dynamics impact the shape of phylogenetic trees. As such, we can expect methods that infer one dynamic aspect while ignoring the other may be biased. To illustrate the nature of the bias, we first use a simple example. We simulated a phylogenetic tree using the exponential coalescent without any population structure. Subsequently, we inferred the effective population sizes, migration rates, and internal node states twice, first assuming constant effective population sizes over time, and then allowing them to grow exponentially. In both cases, we permit for an additional unsampled deme. When not accounting for population dynamics, internal nodes deeper in the tree are inferred to be in another location than the samples (see Figure S2A). The effective population size of that second location is inferred to be much smaller than the first location (see Figure S2B). The smaller effective population size roughly corresponds to the effective population size early during the exponential growth (figures S2E). The backward-in-time migration rates are inferred to be much higher from the sampled into the unsampled location than vice versa (figure S2C). Without correctly accounting for population dynamics, the unstructured exponentially growing population is explained by a small population with strong migration into a larger population. Based on this illustration, we would expect to overestimate the number of introductions from a deme with few into a deme with many samples. When a deme has only a few samples, the effective population size of that deme essentially becomes unconstrained by any data, which the model will use to approximate past population dynamics.

We illustrate this issue using Zika virus (ZIKV) dnd show how accounting for population dynamics can recover more plausible ancestral state reconstructions. We use a previously analyzed dataset of ZIKV sequences sampled from Polynesia, Brazil, the Caribbean, and various locations in South America (Faria *et al*., 2017). This study used DTA to infer that ZIKV was most likely introduced once into the northeast of Brazil, followed by subsequent spread in Brazil and elsewhere in the Americas (Faria *et al*., 2017; Grubaugh *et al*., 2017; Black *et al*., 2019). We perform two different inferences: first, we assume the effective population sizes to be constant over time, and second, we allow them to vary over time. We jointly infer the phylogenetic tree, evolutionary rate, and population parameters under the structured coalescent assuming constant forward-in-time migration rates.

As shown in figure 2A and B, the ancestral state reconstructions vary greatly when accounting for population dynamics (Skyline) and when not (Constant). In the skyline scenario, we infer one introduction from Polynesia to the northeast of Brazil and from there to the other parts of Brazil and the Americas. In the constant scenario, on the other hand, we infer a introduction of ZIKV from Polynesia to the Caribbean and subsequently to different regions in Brazil (see figure 2B and F).

**Figure 2:**
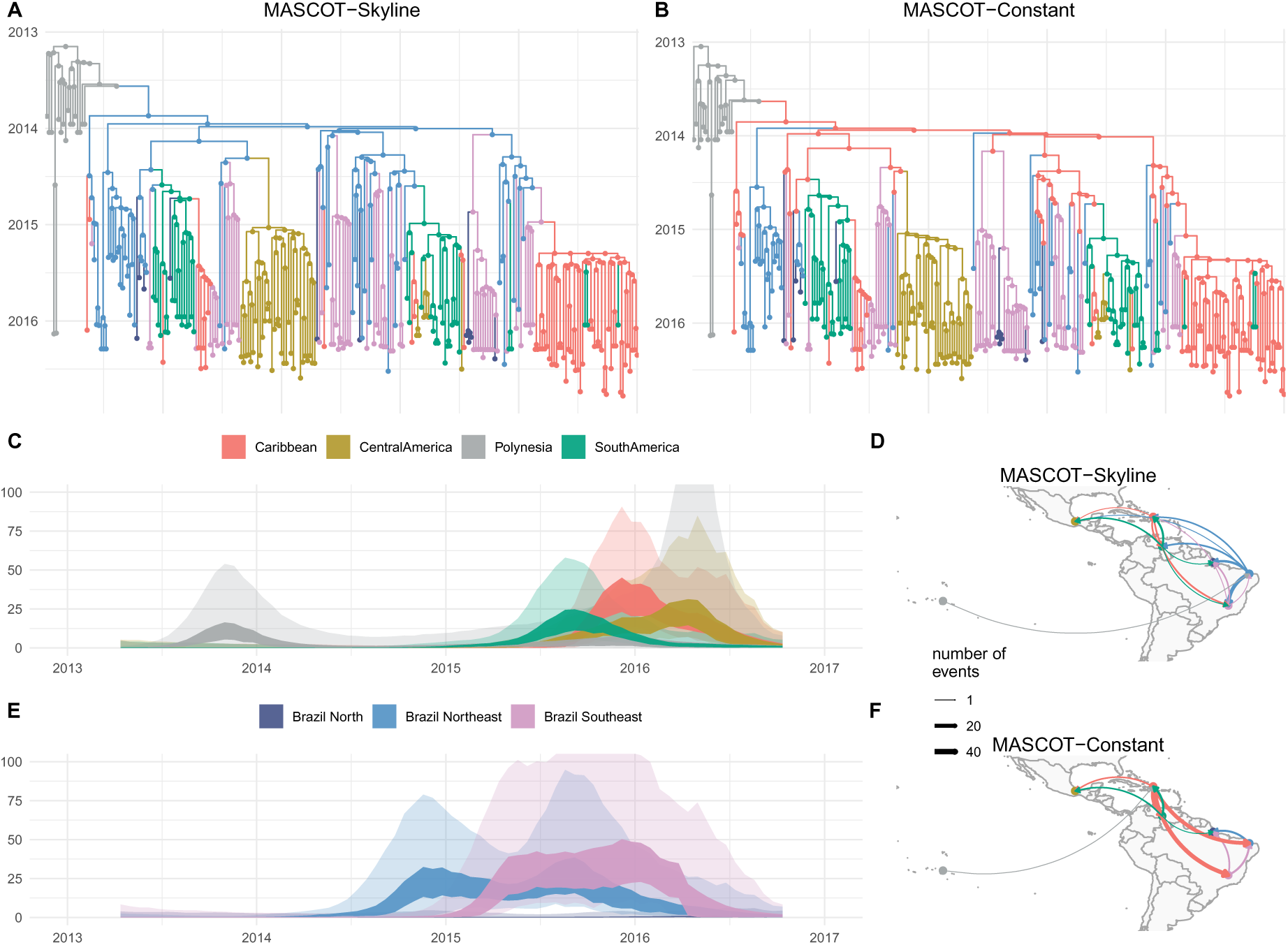
Inferred transmission dynamics of ZIKV when having skyline or constant *Ne* dynamics. **A** Inferred node states when inferring non-parametric skyline Ne’s in different demes. The tree is the maximum clade credibility (MCC) tree, and the nodes are colored by the node with the highest posterior probability in the MCC tree. **B** Inferred node states when each location has a constant Ne over time. **C** & **E** Inferred Ne trajectories for MASCOT-Skyline. The inner interval (dark) denotes the 50% highest posterior density (HPD) interval, and the outer interval (light) the limits of the 95% HPD interval. Inferred number of migration events between the different location using MASCOT-Skyline **D** and MASCOT-Constant **F**.

### Population structure biases population dynamic inference

As previously shown (Heller *et al*., 2013), population structure can impact the inference of population dynamics in coalescent skyline approaches. In particular, reductions in the effective population sizes towards the present can signal sub-population structure that is not accounted for (Heller *et al*., 2013).

To investigate these biases, we compare how the inference of population dynamics is impacted when outside introduction into that population is not accounted for. To do so, we compiled a dataset with influenza A/H3N2 sequences sampled only from New Zealand and Australia, which we denote below as Oceania, sampled between 2000 and 2005. Oceania is thought to mainly act as a sink population for influenza A/H3N2, where there are introductions of viruses into the country that spark annual influenza epidemics, but viruses circulating in Oceania rarely seed epidemics elsewhere in the world (Bedford *et al*., 2010; Bahl *et al*., 2011).

Using this example dataset, we inferred the population dynamics in Oceania twice. First, we assumed no introduction of viruses into Oceania, as well as no export of viruses out of Oceania. We then inferred the effective population size of influenza A/H3N2 into Oceania over time. Next, we allowed for an outside deme to represent influenza transmission anywhere outside of Oceania. This outside deme, sometimes referred to as a ghost deme (Beerli, 2004; Slatkin, 2005), does not have any sampled sequences in the dataset. We estimated the effective population sizes of that outside deme over time alongside the migration rates between Oceania and the outside deme.

As shown in Figure 3, the effective population size estimates are substantially different if we allow for an outside (ghost) deme compared to when we do not allow for that deme. If we allow a ghost deme, the inferred seasonality is much more pronounced. On the other hand, if we do not allow for a ghost deme, we see that the inferred effective population size dynamics of Oceania closely resemble the dynamics of the ghost deme.

**Figure 3:**
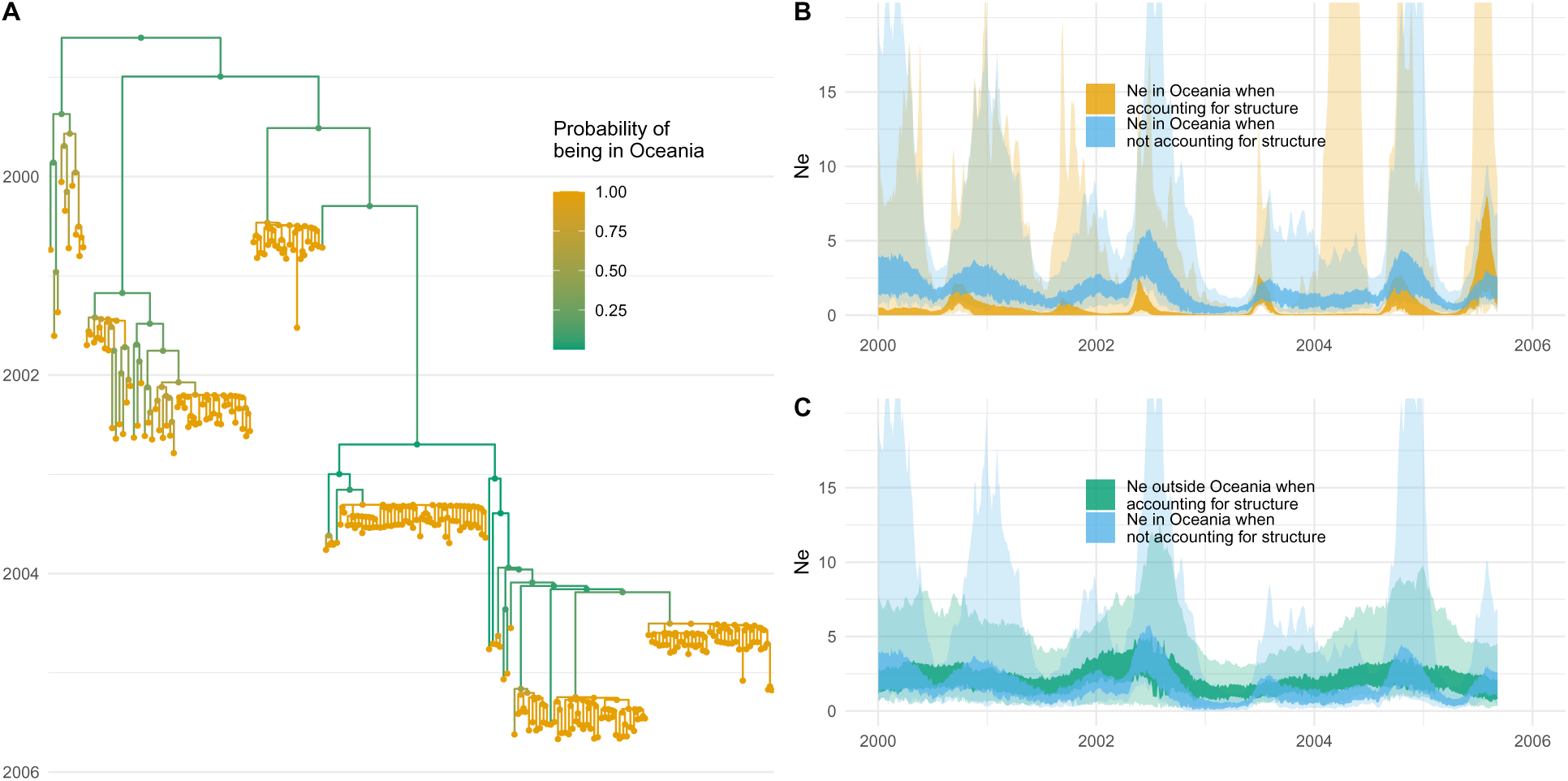
Misinterpretation of population structure as population dynamics for H3N2 in Oceania. **A** Inferred phylogenetic tree of 200 influenza A/H3N2 sequences sampled in New Zealand and Australia (Oceania). **B** Inferred effective population sizes in Oceania when allowing for an unsampled outside (ghost) deme and when assuming no population structure. **C** Inferred effective population sizes in Oceania when not allowing for an unsampled outside (ghost) deme compared to the inferred effective population size of the ghost deme when allowing for population structure.

We next tested using the same simulations as in Figure 1, what effective population size dynamics a skyline method recovers that does not model population structure. As we show in Figure S4, ignoring population structure in these simulations means that the inferred effective population size trajectories closely resemble the larger population.

**Figure 4:**
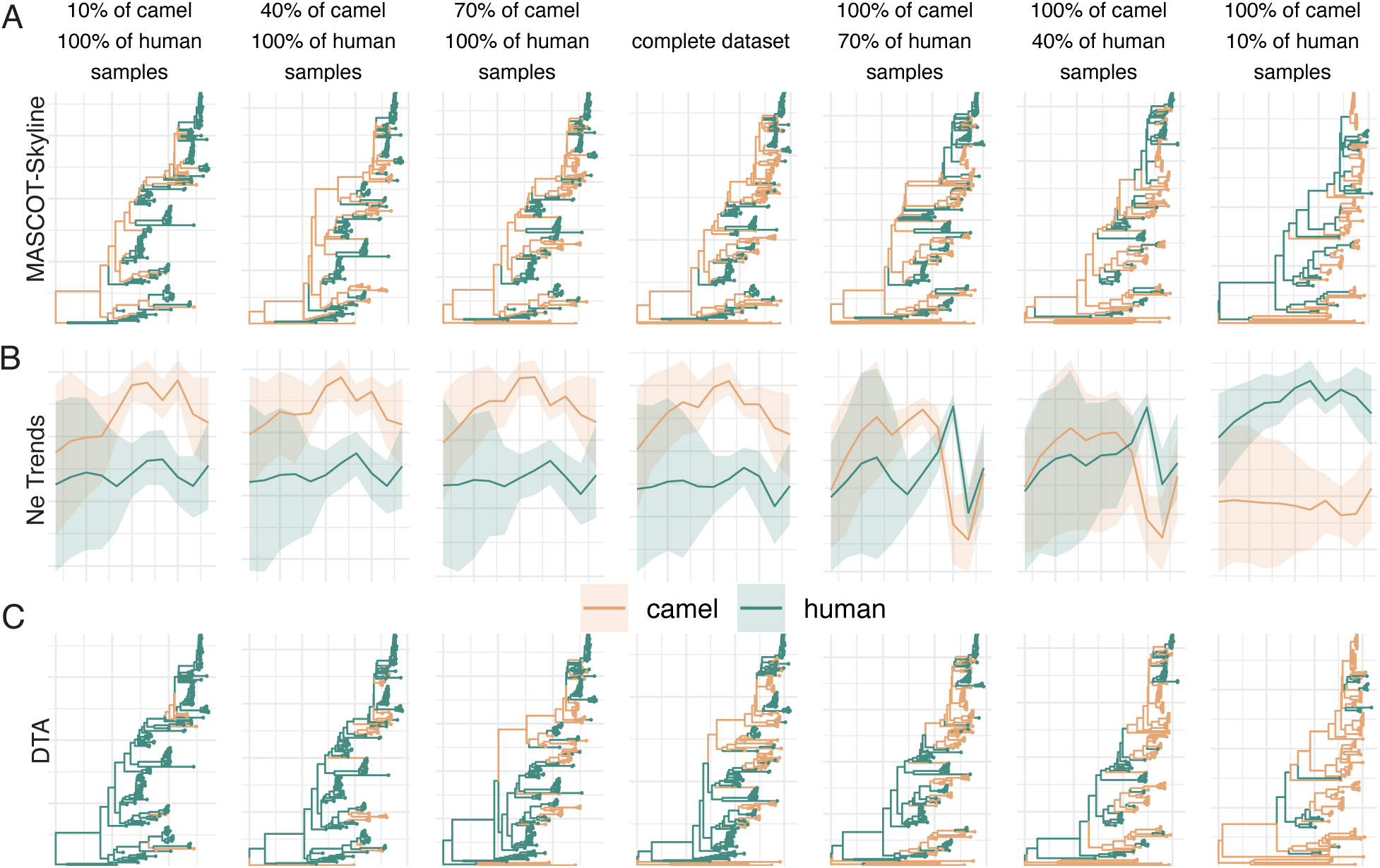
Repeated spillover of MERS-CoV from camels to humans. **A** Maximum clade credibility (MCC) trees inferred using MASCOT-Skyline for different amounts of samples from camels and humans, from left to right). Each branch is colored by the most likely location of the child node of that branch. **B** Inferred effective population size trajectories using MASCOT-Skyline for different amounts of samples from camels and humans. **C** Maximum clade credibility (MCC) trees inferred using DTA.

### Sampling bias impacts ancestral state reconstructions

The coalescent patterns in phylogenetic trees indicate where lineages are over time. For example, rapid coalescence indicates smaller populations. If lineages rapidly coalesce, they are more likely to be in a smaller population.

Here, we investigate the power and pitfalls of this by reconstructing the transmission of MERS-CoV between camels and humans using the dataset from Dudas *et al*. (2018). MERS predominantly circulates in camels with occasional spillovers followed by limited transmission in humans. The dataset described in Dudas *et al*. (2018) contains 274 sequences sampled from humans and camels. We subsampled this dataset ranging from 100% of human samples and 10% of the camel samples to 100% of both and then to 10% of the human samples and 100% of the camel samples. We then performed ancestral sequence reconstruction using MASCOT-Skyline and DTA.

As shown in Figure 4, when there are few camel samples, DTA infers MERS to circulate in humans with occasional spillovers into camels. With all 274 sequences in the data, DTA still infers the predominant circulation in humans. Only when most human samples are removed DTA start to infer the predominant circulation in camels.

Conversely, MASCOT-Skyline infers predominant circulation in camels, even if most camel sequences are removed. The reason for that is that the human samples indicate rapid coalescence and, therefore, a small Ne. For branches that do not conform with a small Ne, it infers them to be in the larger outside (here camel) population. When we remove more and more human sequences, the picture changes. The more recent camel sequences are strongly clustered geographically, also indicating a small Ne. Now that there are fewer human sequences, the human Ne effectively takes the role of a “ghost” deme, and MASCOT-Skyline infers rapid coalescence (that is, the local outbreak clusters) after introductions from elsewhere. Since the only possible location for elsewhere is the human compartment, MASCOT-Skyline infers that local outbreak clusters have been introduced from outside. Interestingly, this means that the biases are inverted between the MASCOT-Skyline and DTA, with MASCOT-Skyline being more likely to infer a human source with fewer human samples.

We next remove local outbreak clusters by first identifying groups of sequences sampled from the same location in the same month. We then only use one of the sequences from that group to represent the outbreak. When we remove local outbreak clusters in the camel compartment, we infer camels to be the source location much more consistently across different sample numbers (see figure S5). We infer circulation in humans only when using almost exclusively camel sequences.

**Figure 5:**
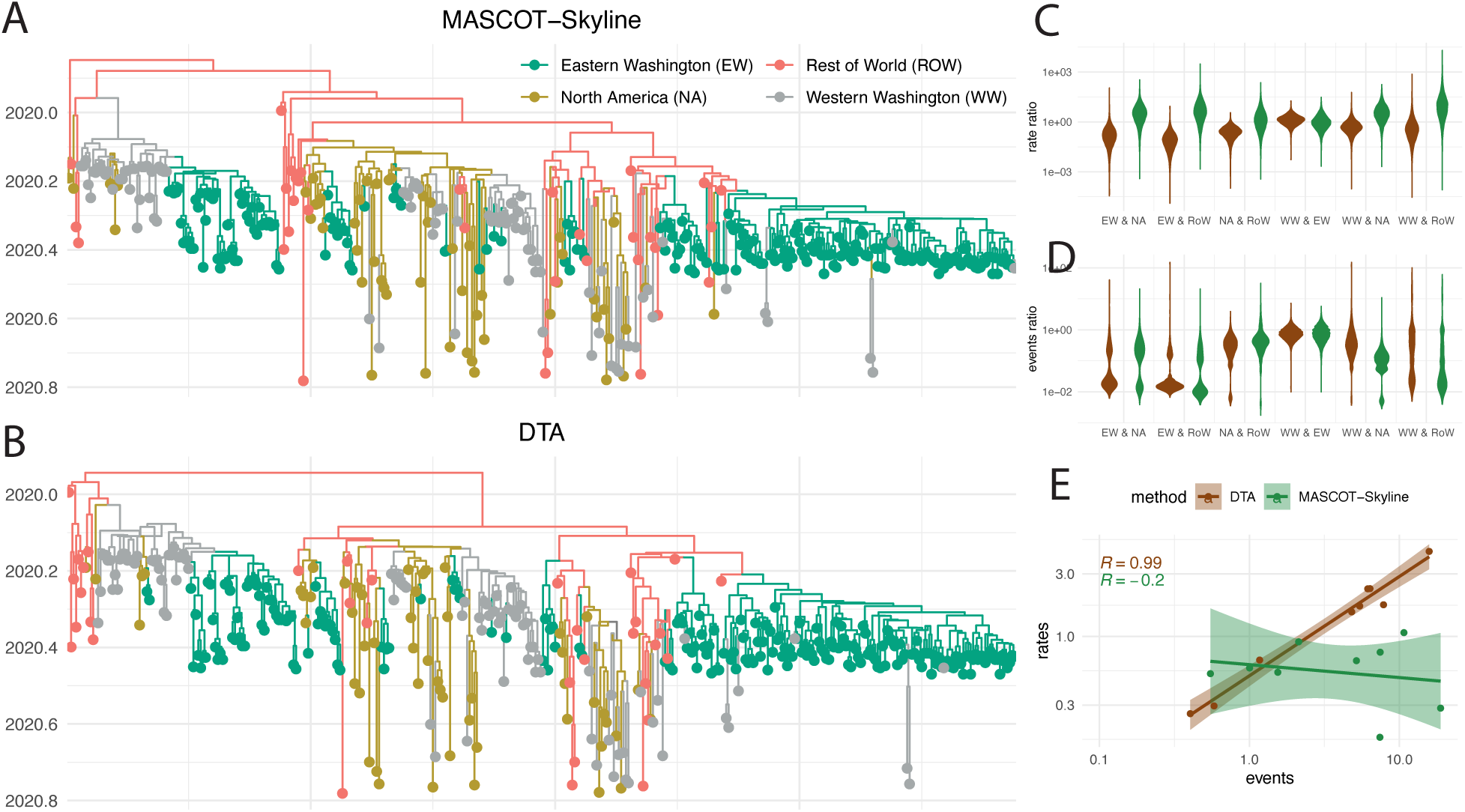
Reconstruction of the geographic spread of SARS-CoV-2 between the world, North America, and Eastern and Western Washington. **A** Maximum clade credibility tree reconstructed using MASCOT-Skyline and DTA (**B**). The colors represent the inferred node states with the highest posterior probability. **C** Inferred migration rate ratios between the four locations using MASCOT-Skyline and DTA. Each violin plot shows the rate ratio from A to B over the rate of B to A. **D** Inferred number of migration events between the four locations using MASCOT-Skyline and DTA. Each violin plot shows the number of migration events from A to B over the number of events from B to A. **E** Correlation between the inferred migration rates and the number of migration events between the four locations. The correlation coefficients are calculated using the median number of events between the 4 locations and the median migration rates between them.

### Modeling population size dynamics is necessary to reconstruct migration rates

When we analyze spatial transmission patterns, we typically seek to infer the movement of viral lineages and/or the rates governing that movement. Reconstructing the movement of viral lineages—performing ancestral state reconstruction—can reveal how many introductions occurred in a location and the number of migration events between locations. However, the number of events identified directly correlates to the number of samples in a location. The more we sample from a location, the more introductions into that location we will identify. The migration rates are population-level parameters independent of the number of samples. The migration rates also tend to be more important to understanding the spread of pathogens than solely the number of migration events.

Importantly, migration rates can be used to determine what drives spatial transmission dynamics, such as using generalized linear models (GLM) (Lemey *et al*., 2014). In the GLM approaches (Lemey *et al*., 2014; Müller *et al*., 2019), the contribution of predictors to the migration rates is inferred instead of directly inferring these rates. Yet, this still relies on the models’ ability to quantify migration rates accurately.

We next show, starting from the example of SARS-CoV-2, how well ancestral states and migration rates can be inferred using MASCOT-Skyline and discrete trait analyses (Lemey *et al*., 2009). We use sequences collected from Washington state (USA), North America, and the rest of the world, previously analyzed in (Müller *et al*., 2021). We further split sequences in Washington state into eastern and western Washington state based on whether the county of isolation is east or west of the Cascade mountain range. We then performed phylogeographic analyses using MASCOT-Skyline and DTA.

As shown in figure 5A and B, DTA and MASCOT-Skyline infer similar ancestral state reconstructions. These similar ancestral state reconstructions reflect similar migration events between the four discrete locations (Fig. 5D). To further quantify the similarity in the ancestral state reconstructions between the two methods, we infer the sampling location of 5 random tips from each location that have had their location masked before running phylogeographic inference. We then computed the posterior support of the sampled location to be in the correct location of isolation. As shown in figures S6, the posterior support for the correct location of isolation is similar between the two methods. However, DTA has a higher posterior support for the actual sampled location than MASCOT-Skyline.

While the two methods reconstruct similar ancestral states, they infer vastly different migration rates (Figure 5C). In particular, DTA infers migration rates highly correlated to the number of migration events between two locations (Figure 5E). The migration rates inferred by MASCOT-Skyline instead have little to no correlation with the number of migration events between the two locations. If we have two locations, one with ten times more number of infected individuals, then we would expect ten times more migration events from that location, even if the migration rates are the same. Therefore, this means that the number of migration events is not a sufficient measure of the migration rates. Since DTA does not incorporate population dynamics into the estimation of migration rates, these differences are not unexpected.

Unlike the sampling location, we do not know the actual migration rates for this dataset. However, we can use simulations to investigate when the two methods perform well. To this end, we perform simulations using a Susceptible-Infected-Recovered (SIR) model with two states using MASTER (Vaughan and Drummond, 2013). We perform SIR simulations using various sampling models, different migration rates, and different reproduction numbers R0 across states.

MASCOT-Skyline is able to recover the prevalences over time for the two states (see figure S7 & S8). Both methods, DTA and MASCOT-Skyline, can recover ancestral states similarly well for low rates of migration (see figure S9). DTA has greater posterior support for both the right and the wrong node states (see figure S9). Overall, both approaches recover the true ancestral node states similarly well, which is consistent with our analyses of the SARS-CoV-2 dataset.

As suggested by our SARS-CoV2 analyses, we find large differences in the migration rate estimates between the two methods (see figure S10). MASCOT-Skyline recovers the rates accurately for most simulation scenarios, with somewhat worse performance when R0’s differ across the two states (see figure S10). This was expected based on our assumption that the prevalence is proportional to the effective population size with the same proportionality factor across states. We therefore expect that explicitly accounting for differences in these proportionality factors would remedy these biases. DTA overall suffers from relatively low coverage of the true value in these simulations of between 27% and 89%. These low coverage values are partly explained by a lower correlation between true and estimated values but also by narrower highest posterior density intervals (see figure S11). Both methods are able to retrieve the magnitude of migration, that is, the mean migration rate accurately (see figure S12). The estimated mean migration rates are highly correlated to the simulated values, though DTA has lower coverage of the true simulated values due to narrower HPD intervals. Lastly, we compared the ratio of migration rates from state 1 to 2 over the migration rate from 2 to 1.

Lastly, we investigate if correcting for the cumulative prevalence in the source and sink locations for DTA improves the correlation of the migration rate estimates. We find some improvement, but the correlation is still weaker than for MASCOT-Skyline (see figure S10).

## Discussion

Here, we show that population dynamics and population structure are intrinsically linked when inferring the spread of pathogens. This is consistent with previous work on biases in phylogeographic (Layan *et al*., 2023) and phylodynamic models (Heller *et al*., 2013). To address this, we develop MASCOT-Skyline, an approach to infer non-parametric population dynamics alongside population structure.

Using the example of ZIKV spread in South America, we show that assuming the wrong population dynamic model dramatically impacts the reconstruction of how the spread of ZIKV unfolded, with MASCOT-Skyline providing a reconstruction that is much more consistent with other estimates (Faria *et al*., 2017).

The bias introduced by assuming constant effective population sizes over time is relatively hard to predict *a priori*. Anecdotally, locations with very few samples can act similarly to a ghost deme (Beerli, 2004; Slatkin, 2005). In that case, the Ne of locations with only a few samples is potentially used by the model to approximate the population dynamics of the other state. While we did not explicitly investigate performance differences between MASCOT-Skyline and constant, the computational demands for MASCOT-Skyline do not seem to be substantially higher than for the constant approach. This is particularly true when the Ne is only estimated at, for example, ten or fewer time points. Population dynamics are present to some degree in most datasets, which should make approaches that account for them better suited to analyze these datasets in all but a few cases. As such, we recommend defaulting to MASCOT-Skyline over constant.

Using the example of MERS-CoV, we illustrate this bias by changing the number of samples from humans and camels. MERS-CoV circulates in camels and repeatedly spills over into humans, causing limited outbreaks. If the sink population is extremely undersampled, the effective population size of the sink population will be used to approximate the population dynamics of the source population. Interestingly, this leads to the opposite sampling bias than in the case of discrete trait analyses (DTA) (Lemey *et al*., 2009). DTA tends to assign the source location to the overrepresented deme. The fewer human samples there are, the more likely camels are inferred to be the source location. This is caused by the sampled numbers being treated as informative by DTA. While the explanation for the pattern inferred by DTA is relatively straightforward, the explanation for the pattern inferred by MASCOT-Skyline is more complex. We suspect that with more camel samples and only a few human samples, MASCOT-Skyline infers a Ne trajectory for the camel state that is consistent with the local outbreak clusters. This opens interesting questions about what level of structure is important to consider in such analyses and how to choose samples that reflect that level of population structure. In the case of MERS-CoV, if one is interested in the structure at the level of the host species, sampling (or subsampling) has to be performed to represent this structure. As such, doing so requires information about the sampling process and the potential exclusion of some of the sequences collected, for example, from outbreak clusters.

Ancestral state reconstruction provides a picture of the path of individual lineages. Additionally, ancestral state reconstructions can act as a sanity check on whether the inference results are consistent with prior knowledge, such as which species is the host species. Using the example of SARS-CoV-2, we show that similar ancestral state reconstructions can lead to vastly different migration rate estimates between DTA and MASCOT-Skyline. While we do not know the true migration rates in this case, the rate estimates of MASCOT-Skyline are more consistent with what is expected from the population sizes of the different locations in the dataset.

Based on our simulation study and the SARS-CoV-2 example, the migration rate estimates by DTA should likely not be interpreted as population-level parameters in most cases. That is, they do not reflect the rate at which an individual in location A move to location B, unless the sampling rates are constant over time and the same across locations A and B. Instead, they should likely be interpreted as a parameter that mainly reflects the observed number of migration events between locations A and B in the dataset. Therefore, subsampling strategies for DTA analyses should likely be based on the number of infected over time and across locations. If that is not possible, the migration rate estimates may not be directly interpretable as epidemiological parameters. This also poses interesting questions for methods that seek to reconstruct the drivers of migration patterns, for example, using generalized linear models (Lemey *et al*., 2014; Müller *et al*., 2019). The results of such analyses could also be subject to similar limitations.

Sampling biases are a persistent challenge to phylogeographic reconstructions. This research shows that it is crucial to consider the sampling process in phylogeographic reconstructions for relatively simple (DTA) and more complex models (structured coalescent). Further, we show that migration dynamics must be considered in a population dynamic context.

## Methods and Materials

### MASCOT

MASCOT, the marginal approximation of the structured coalescent, tracks the probability of lineages being in any of the modeled states backward in time by solving ordinary differential equations described in Müller *et al*. (2017) and Müller *et al*. (2018). MASCOT is parameterized by effective population sizes and migration rates. The effective population size of state a is given by *Ne_a_*, and the backward migration rate from state *a* to state *b* is given by *m_ab_*. MASCOT assumes that the effective population sizes and migration rates are constant during each integration step. By solving the ordinary differential equations (ODE), MASCOT computes the probability of the tree given the parameters 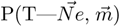. To model time-varying parameters, we feed the continuously varying values for 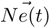 and 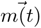 into the ODE calculations as piecewise constant values 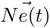 and 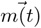 at different time points t that approximate the underlying continuous dynamics. The piecewise constant approximation uses a user-defined number of intervals, with more intervals leading to a better approximation of the continuous dynamics of the parameters but also higher computational costs. We further explain this in figure S14. The probability 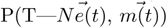 can then be computed by integrating over all possible states at the root of the tree Müller *et al*. (2018). Additionally, one can compute the probability of each node in the tree being in any state to perform ancestral state reconstruction (Müller *et al*., 2018) or explicitly reconstruct the migration histories using stochastic mapping (Stolz *et al*., 2022).

### MASCOT-Skyline

To model nonparametric population dynamics alongside population structure, we first define a grid of time points to model nonparametric population dynamics. We define the grid in absolute time or relative to a tree’s height, which is the default option. We then infer each grid point’s effective population size 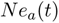. Between those points, we assume that the effective population sizes change continuously according to an exponential growth model. Effectively, we use linear interpolation between any two adjacent Ne’s in log space. Alternative approaches, such as spline interpolation, would also be possible to implement. For the computation of 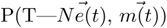, we approximate the continuous parameter dynamics using piecewise constant approximation as described above and then use the piecewise constant values for the integration of the MASCOT ODE’s. Typically, the number of intervals used for the piecewise constant approximation should be substantially higher than the number of the Ne’s estimated for this to be a reasonable approximation.

By default, we assume the forward-in-time migration rates to be constant over time. As the backward-in-time migration rates that go into the computation of P(T—*θ*), we say that the backward-in-time migration rate 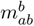 from *a* to *b* is:

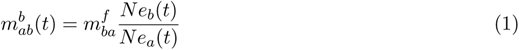

Using the derivation of the coalescent rates or effective population sizes in (Volz, 2012), the error *ɛ* of this assumption is:

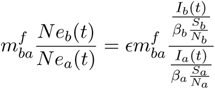

And, therefore

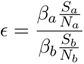

meaning the error we introduce equals the effective transmission rate in the sink divided by the rate over the sink. With the reduction of the pool of susceptible individuals *S_a_* in the population *N_a_*, the difference induced by differences in the transmission rates *β* will likely become smaller. Therefore, the error of the assumption that the ratio of Ne’s between source and sink is equal to the ratio between the number of infected individuals is reduced over time. However, in cases where there is a difference in, for example, the generation time (or the becoming uninfectious rate), the error will persist. For example, this could be the case when studying the transmission across different host species.

In addition to the skyline model, we implemented exponential and logistic growth models. The different dynamic models for the effective population size can be mixed. For example, state *a* can be a skyline model, while state *b* can grow exponentially or be constant. The above equation assumes that the ratio of Ne’s between source and sink locations is equal to the ratio in the number of infected individuals.

### Joint inference of effective population sizes and migration rates

To infer the effective population sizes, the different demes, and the migration rates, we use the adaptable variance multivariate normal operator Baele *et al*. (2017). The adaptable variance multivariate normal operator proposes new parameter states during the MCMC and learns the correlation structure between the different parameters. The effective population sizes are denoted in log space, while the migration rates are logged in real space, that is, not in log space. The prior on the effective population sizes, sometimes referred to as a smoothing prior, is similar to the skyline method (Gill *et al*., 2013). The implementation of the smoothing prior works is as follows. One can choose an arbitrary prior on the difference between two adjacent Ne’s in log space. Further, one can choose a prior distribution on the most recent/present Ne. If the prior on the difference between two adjacent Ne’s is a normal distribution with mean 0 and standard deviation σ, then the smoothing prior is a Gaussian Markov random field (GMFR). The σ parameter itself can be fixed or estimated from the data, which corresponds to the precision of the skyline method (Gill *et al*., 2013). By default, selecting the σ to be estimated for each state will mean a different value for σ will be estimated individually for each state. To change between source and sink locations throughout the MCMC, we use an operator that swaps the effective population sizes for the same time points between locations *a* and *b*. All other operators used for the MCMC are the default operators in BEAST2 (Bouckaert *et al*., 2019).

### Implementation

We implemented MASCOT-Skyline as part of the BEAST2 package MASCOT. MASCOT-Skyline requires at least the BEAST2 version 2.7 to execute. The code is available at https://github.com/nicfel/Mascot and through the BEAST2 package manager. MASCOT-Skyline is implemented in Java. MASCOT-Skyline is available starting from MASCOT version v3.0.5. Analyses can be set up using the BEAUti interface of BEAST2 by choosing MASCOT-Skyline as a tree prior. The effective population size dynamics are chosen separately for each location, deme, or state. Therefore, constant, exponential, or skyline effective population size dynamics can all be used in the same analyses, albeit for different states. For setting the specifications of the Gaussian Markov Random Field (GMRF) prior on the skyline dynamics, one has to specify the prior on the difference between adjacent Ne’s (that is, between the Ne at time t and at time t+1) to a normal distribution with mean 0 and standard deviation s. The standard deviation can then be specified or estimated. The standard deviation is estimated, by default, individually for each state. Throughout this paper, we assume that each state’s standard deviation is the same. Implementing MASCOT-Skyline as an open-source packaged to BEAST2 allows users to use the variety of evolutionary models and data sources implemented in BEAST2 or packages to BEAST2, including relaxed clock models or amino acids alignments.

### SIR simulation study

We use a two-state model to aid the interpretability of the results. We simulate outbreaks in two states, each with an R0 of 1.5, a recovery rate of 52, and a random total population size sampled from a uniform distribution between 500 and 10000. The migration rates are sampled from an exponential distribution with a mean of 5 (low migration rate scenario) or 25 (high migration rate scenario). We simulate phylogenetic trees using the SIR model in MASTER (Vaughan and Drummond, 2013). We then use either 250 or 500 samples per state for inference from the phylogenetic trees. Or use a constant sampling rate, conditioning on at least 50 samples per state. On average, the simulations had 389 (low migration) and 431 (high migration) tips. In the constant sampling scenario, we simulated trees with 4000 tips per state. We then subsampled the tips to have 250 samples per state, sampled evenly across time. Importantly, the samples per state will potentially impose implicit constraints on the possible values for other simulation parameters, such as a state’s population size. We performed discrete trait analyses (DTA) using the BEAST v1.10.4 (Suchard *et al*., 2018; Drummond and Rambaut, 2007). For all analyses, we use a coalescent skygrid tree prior (Gill *et al*., 2013). We estimate the mean migration rate and the relative migration rates between locations. We use an exponential prior on the mean migration rate. We use either 5 (low migration rate scenario) or 25 (high migration rate scenario) for the mean of the exponential prior. We use an exponential prior with a mean of 1 for the relative migration rates. This parameterization of the migration rates is necessitated by the parameterization of DTA likelihood calculation, which normalizes relative migration rates.

Next, we infer the migration rates and the effective population size dynamics using MASCOT-Skyline. We use a Gaussian Markov Random Field (GMRF) smoothing prior to the Ne’s over time and estimate the variance. We estimate the Ne at 26 points in time. For the migration rates, we use an exponential distribution with the mean equal to the mean migration rates in the simulations, i.e., 5 or 25.

### Software

All other plots are done in R using ggplot2 (Wickham, 2016), ggtree (Yu *et al*., 2017), and ggpubr (Kassambara, 2018). Convergence is assessed using conda (Plummer *et al*., 2006).

## 0.1 Data

The source code for the MASCOT package, which includes MASCOT-Skyline, is available at https://github.com/nicfel/Mascot and through the BEAST2 package manager. We provide a simple tutorial to help users start with MASCOT-Skyline here https://github.com/nicfel/MascotSkyline-Tutorial. The scripts to set up analyses and plot the results in this manuscript are available from https://github.com/nicfel/MascotSkyline-Material.

## Acknowledgments

N.F.M. is supported in part by NIH NIGMS R35 GM119774 and a Noyce initiative award. T.B. is a Howard Hughes Medical Institute Investigator. The SARS-CoV-2 analyses use sequences submitted to gisaid.org and are downloaded through the GISAID EpiCoV database. We acknowledge the authors for originating and submitting laboratories of the sequences from GISAID’s EpiFlu database, on which part of this research is based. A full list of sequences used can be found here.

## Supplementary material

**Figure S1:**
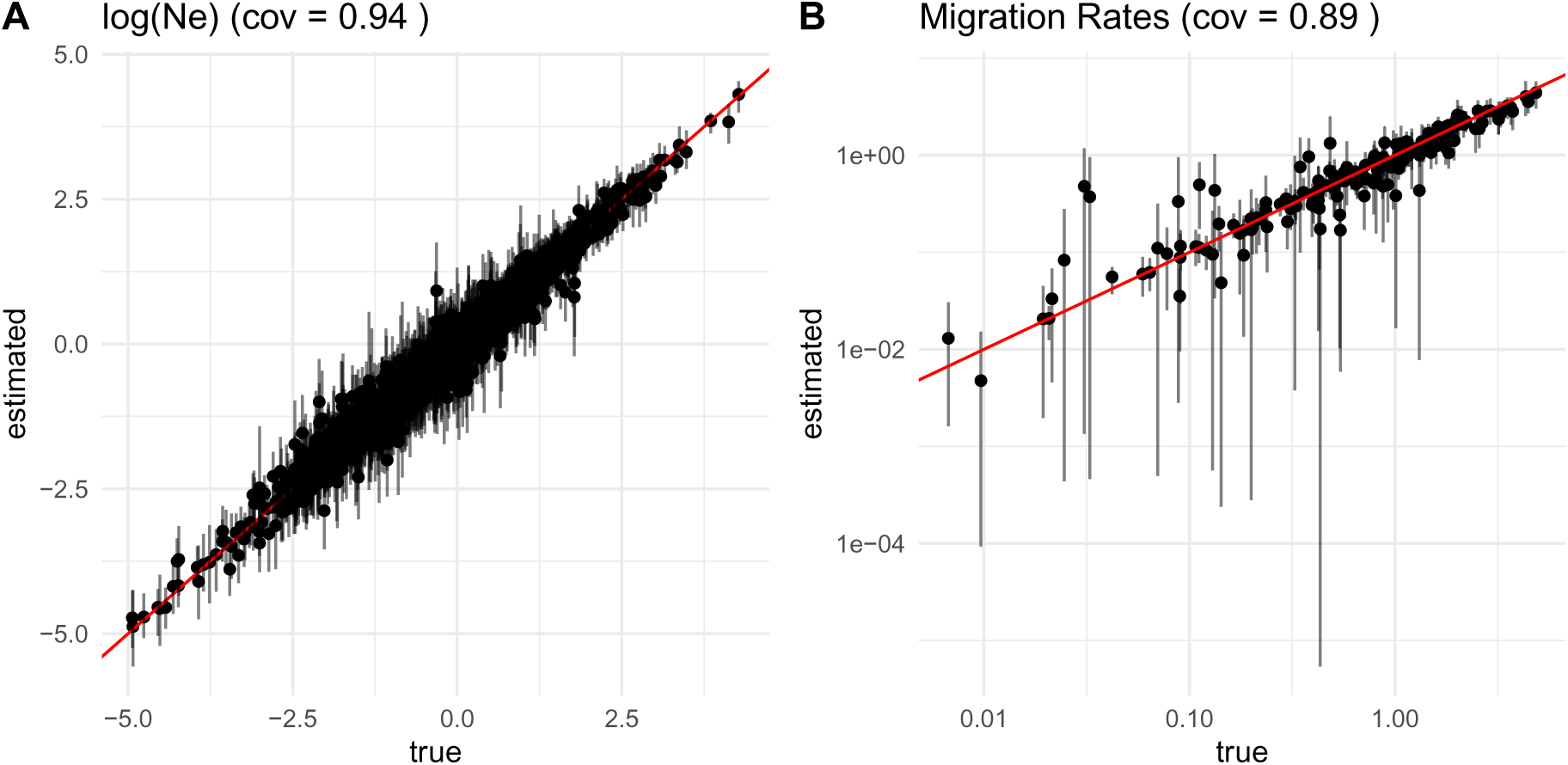
Inferred vs. true effective population sizes and forward in time migration rates for non-parametric *Ne* dynamics. **A** Inferred vs. true effective population size estimates. **B** Inferred vs. true forward in time migration rates. The coverage (cov) of the true value by the 95% highest posterior density interval is shown on the top.

**Figure S2:**
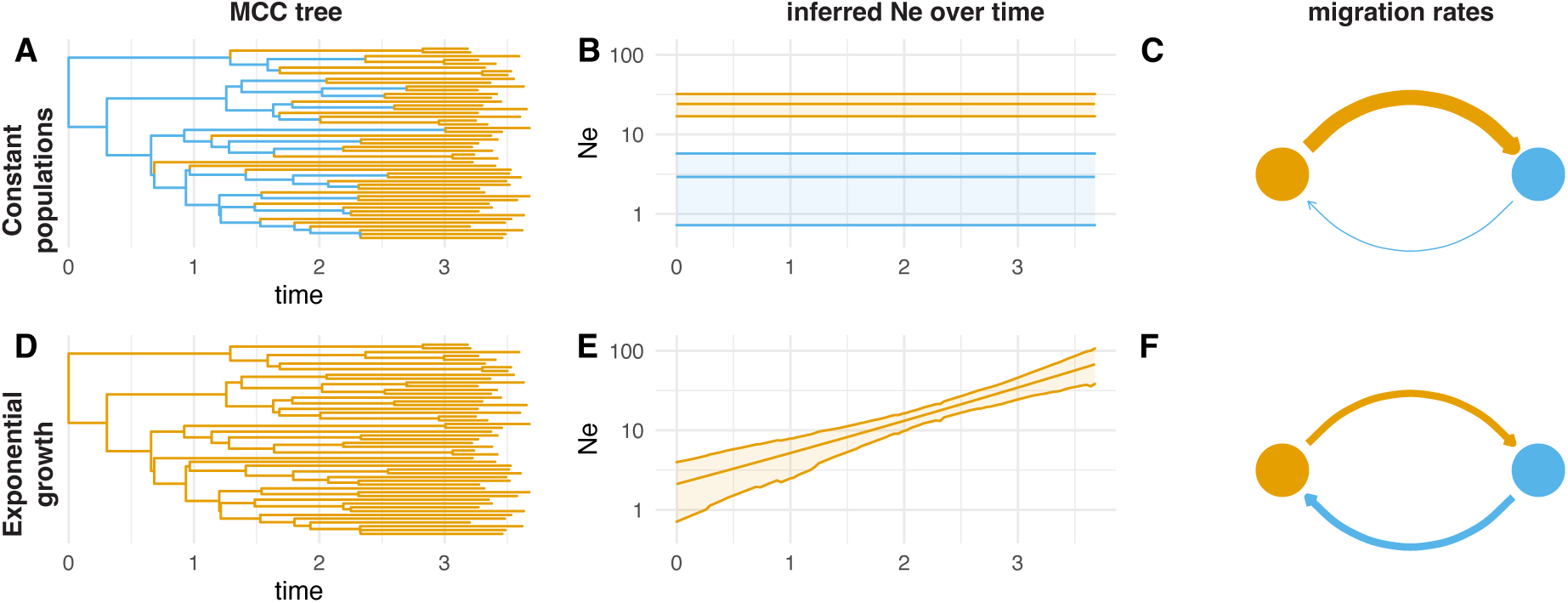
Miss-interpretation of population dynamics as population structure. **A** Inferred node states when assuming a two-state structured coalescent model with two constant populations. **B** Inferred effective population sizes of the two populations. **C** Inferred migration rates between the two constant populations. **D** Inferred node states when assuming a two-state structured coalescent model, allowing the two states to grow exponentially. **E** Inferred effective population sizes over time of the location where all samples were taken from (orange). The Ne of the blue location is sampled under the prior and therefore not shown in the figure. **F** Migration rates between the location where samples were taken and a second (blue) location.

**Figure S3:**
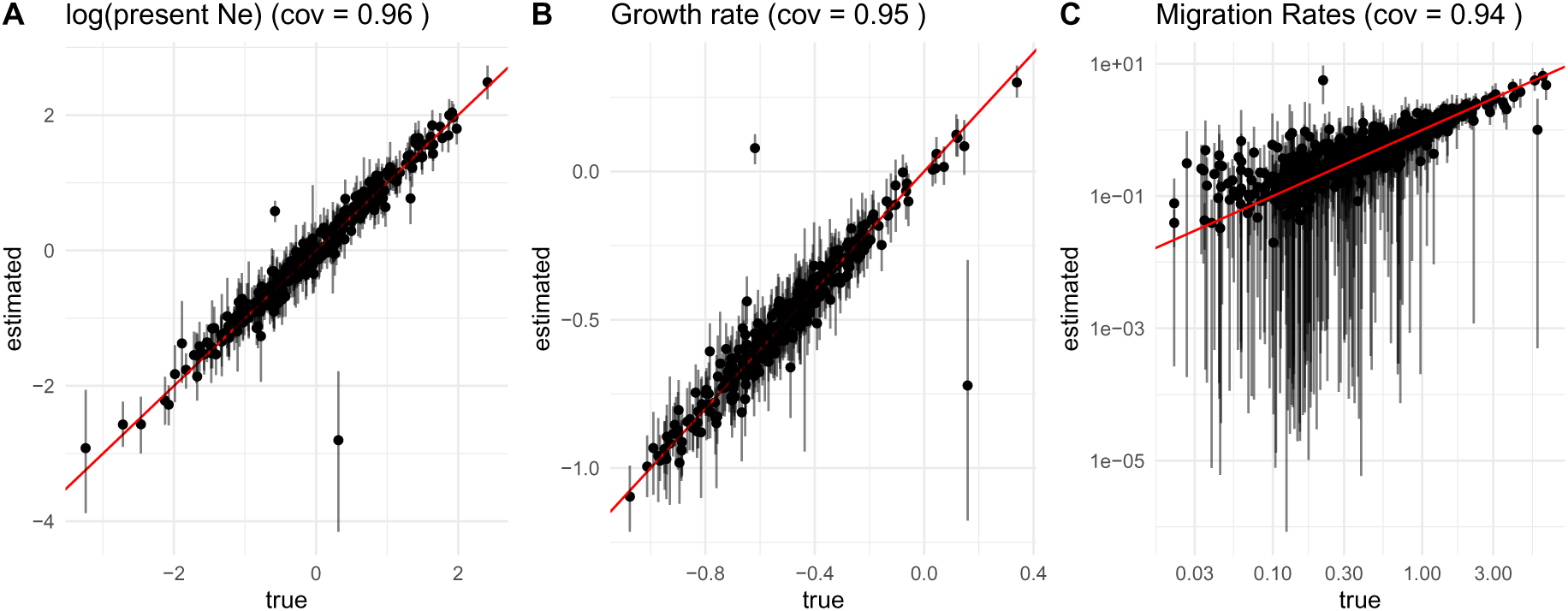
Parameter inference for three states with exponential growth. **A** Inferred vs. true effective log population size at the present. **B** Inferred vs. true growth rates. **C** Inferred vs. true forward in time migration rates. The coverage (cov) denotes how often the true, simulated value was part of the 95% highest posterior density intervals.

**Figure S4:**
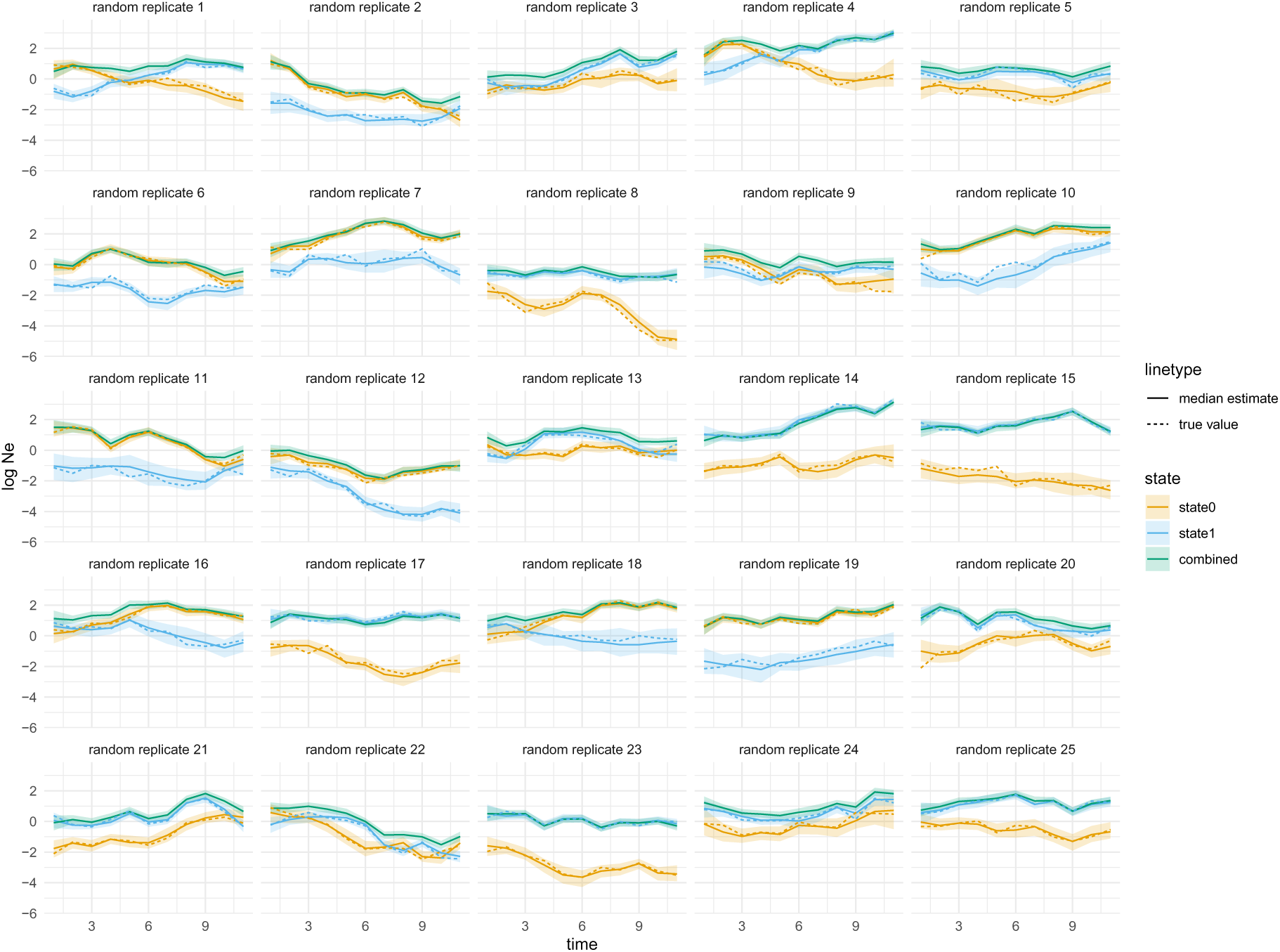
Inferred *Ne* trajectories for a two-state structured coalescent model when population structure is ignored. Here, we infer the effective population size (Ne) trajectory for tree simulation under a two-state structured coalescent model with time-varying population size. We do so once modeling the two states (state 0 in orange and state 1 in blue) and once ignoring any population structure (combined in green).

**Figure S5:**
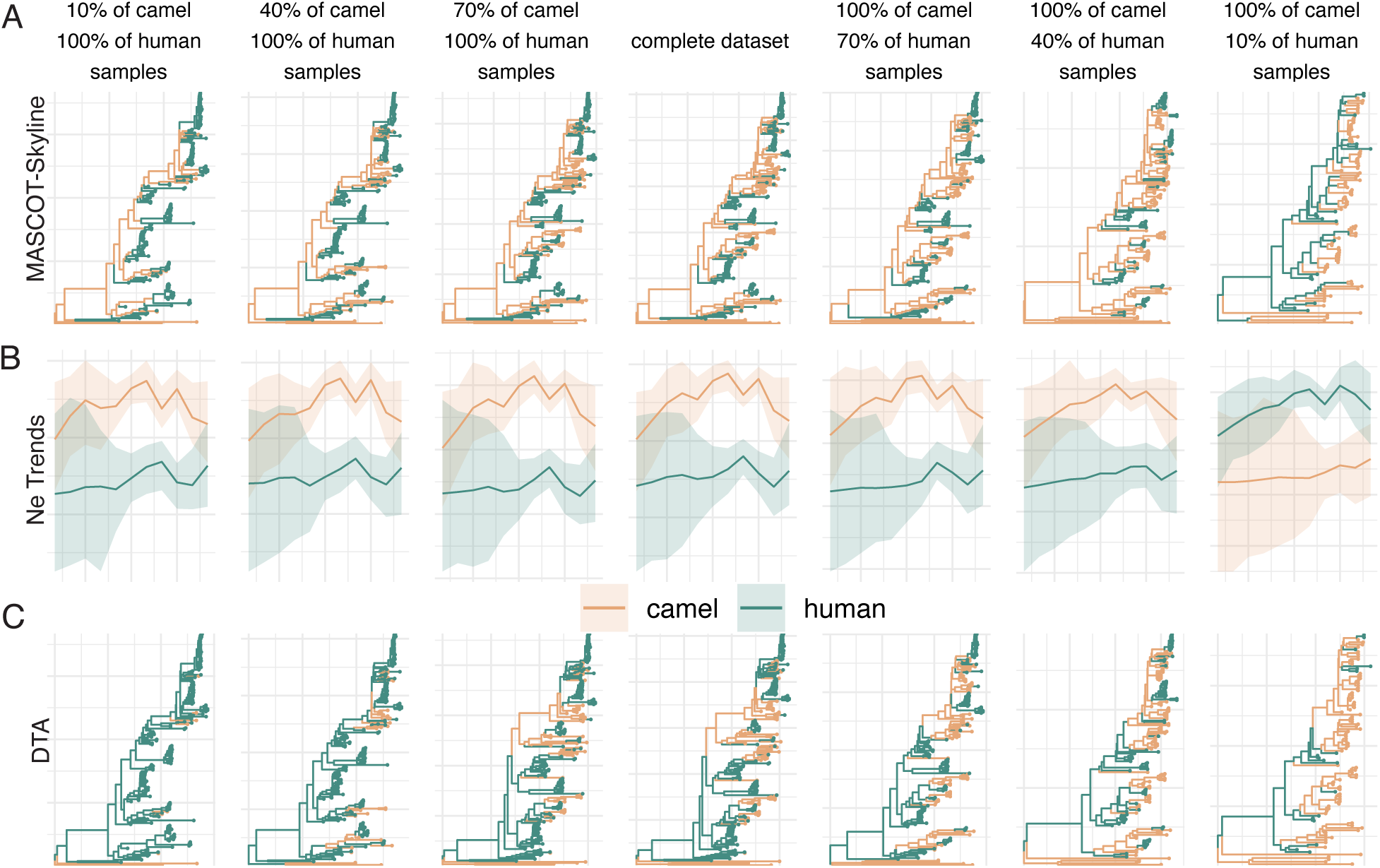
Repeated spillover of MERS-CoV from camels to humans when removing local outbreak clusters. Here, we show the inferences of the transmission dynamics of MERS-CoV between humans and camels when we remove local outbreak clusters in the camel compartment defined as sequences sampled from the same location in the same month. **A** Maximum clade credibility (MCC) trees inferred using MASCOT-Skyline for different amounts of samples from camels and humans, from left to right). Each branch is colored by the most likely location of the child node of that branch. **B** Inferred effective population size trajectories using MASCOT-Skyline for different amounts of samples from camels and humans. **C** Maximum clade credibility (MCC) trees inferred using DTA.

**Figure S6:**
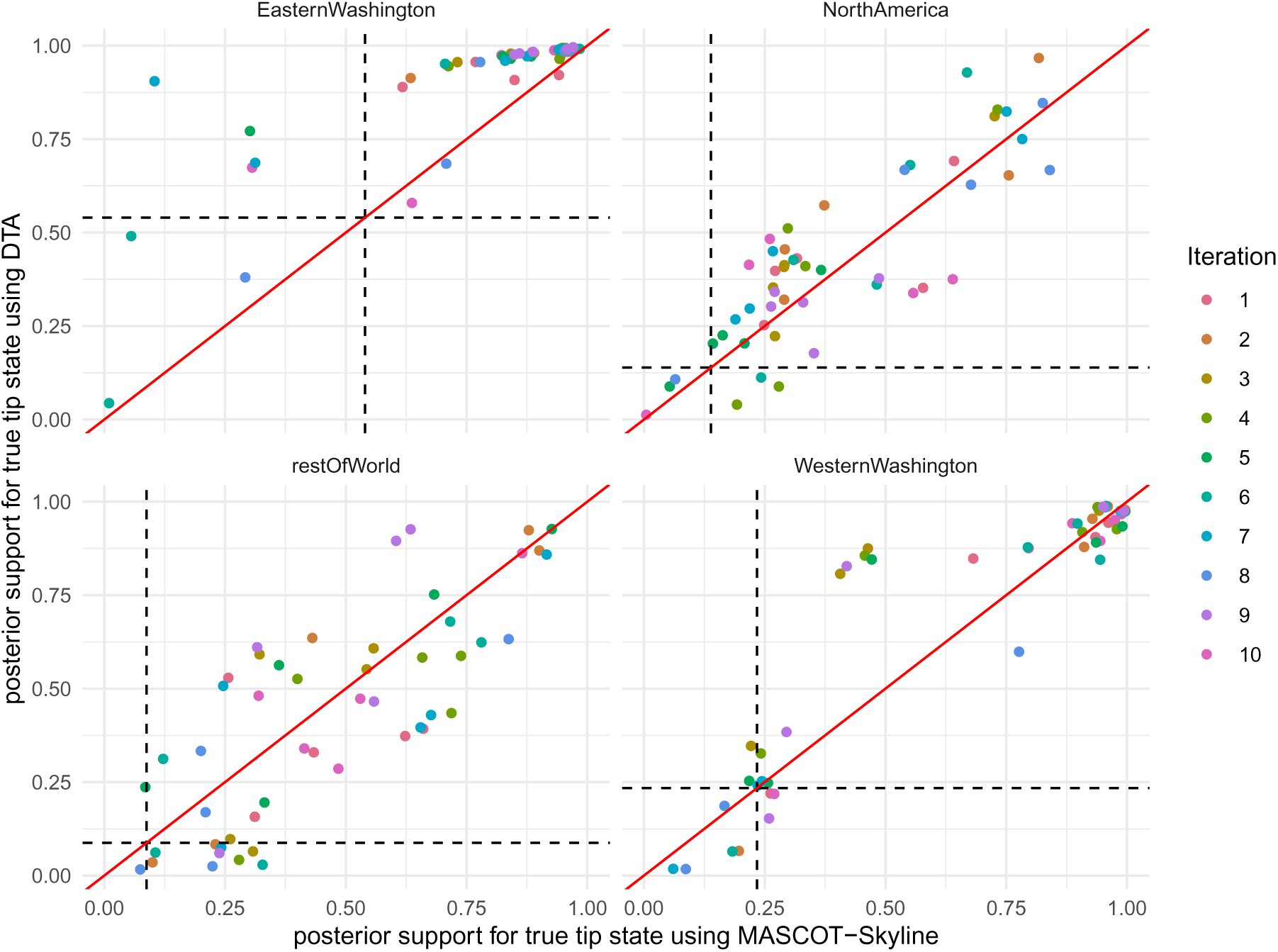
Posterior support for true tip state between MASCOT-Skyline and DTA. We compare the posterior support for the true sampling location inferred using MASCOT-Skyline and DTA for the four locations in our SARS-CoV-2 dataset. For the inference, the sampling location of random samples in the dataset was masked, and the location was re-inferred. The posterior support for the true location then denotes how much posterior weight the MCMC algorithm is putting on the inferred sampling location between the true sampling location. The dotted lines denote the percentage of samples from each geographic location that is in the analyses, i.e., a line at 0.25 would indicate that 25% of samples in the dataset are from that location.

**Figure S7:**
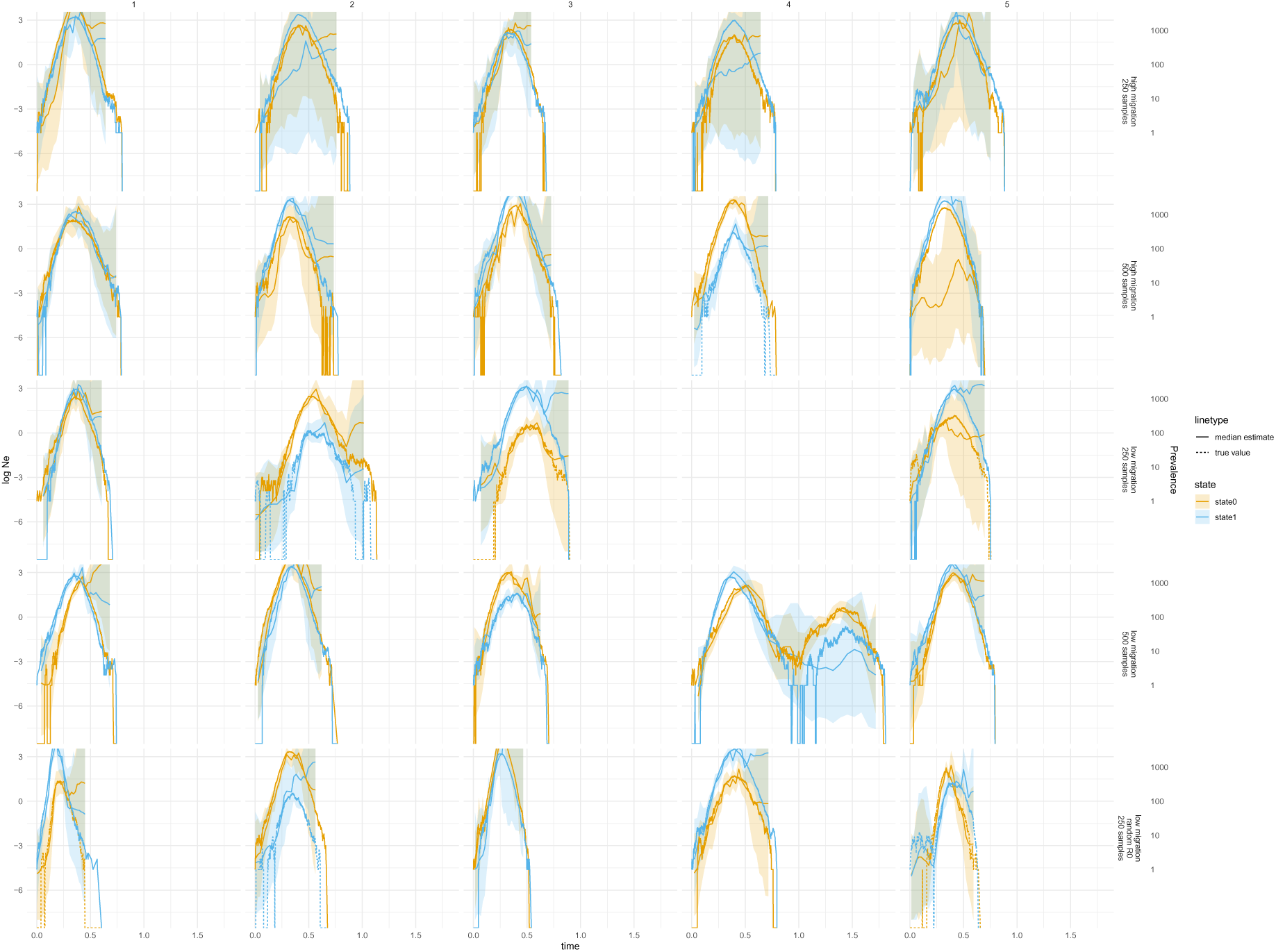
Simulated Prevalence for the two states in the SIR model, part 1. Comparison between simulated prevalences for the two states and the inferred log Ne’s for the two states using MASCOT-Skyline. The trajectories are shown for the first 5 runs of the simulation scenarios denoted on the left.

**Figure S8:**
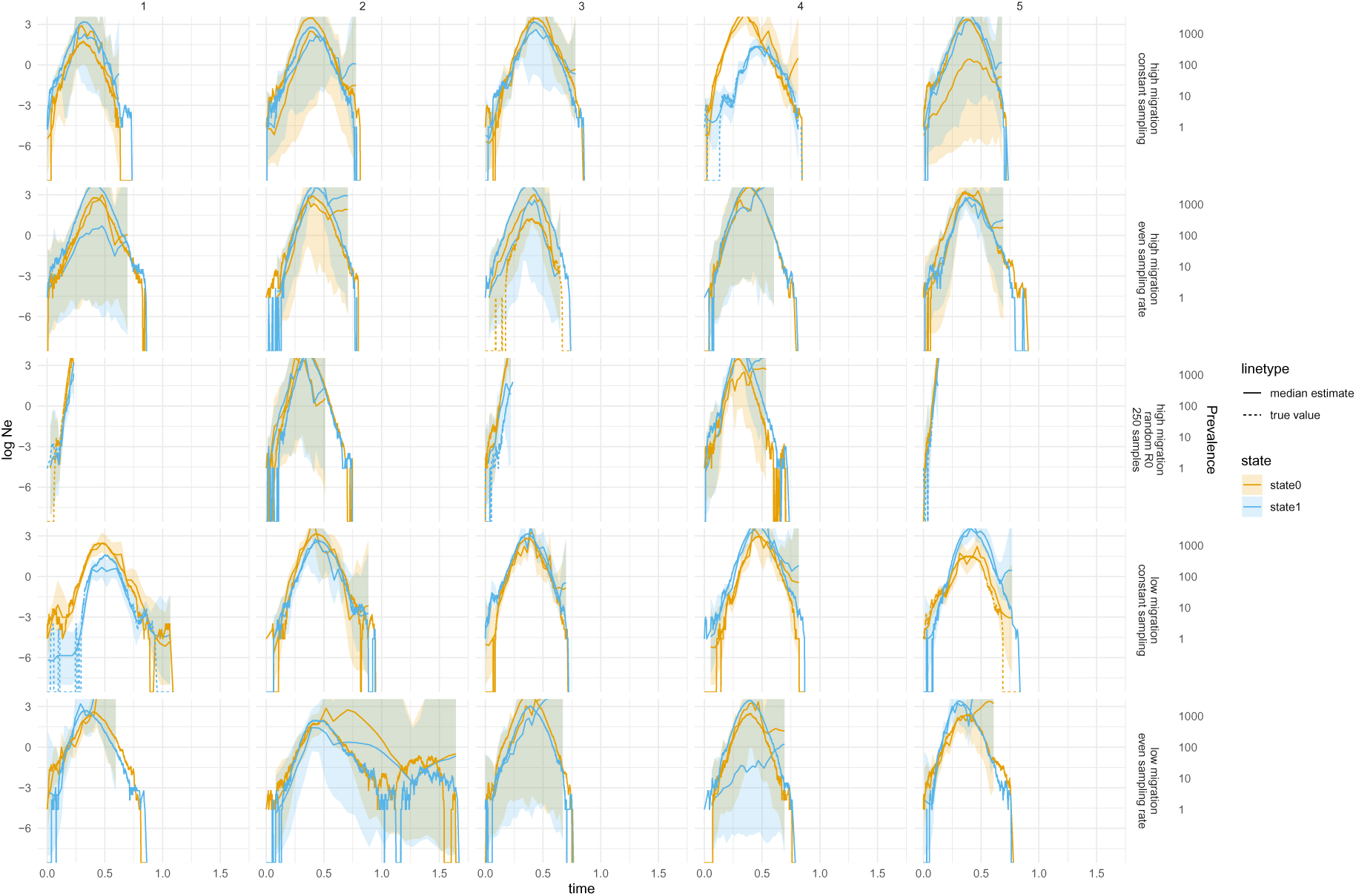
Simulated Prevalence for the two states in the SIR model, part 2. Comparison between simulated prevalences for the two states and the inferred log Ne’s for the two states using MASCOT-Skyline. The trajectories are shown for the first 5 runs of the simulation scenarios denoted on the left.

**Figure S9:**
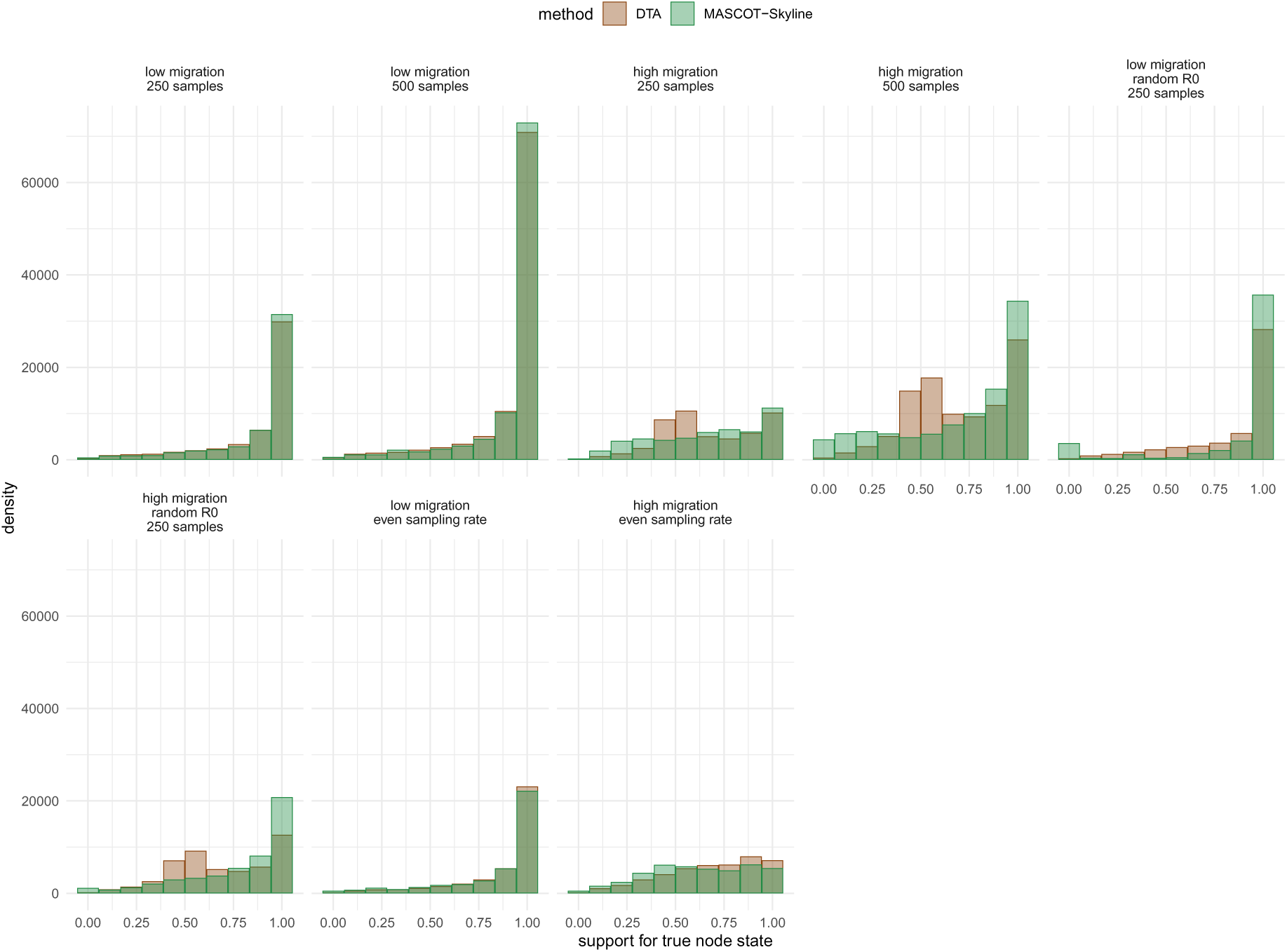
Distribution of the posterior support for the true node states inferred by DTA and MASCOT-Skyline. Here, we show the distribution of posterior support for the true node states for the different SIR simulation settings. The posterior node supports are shown DTA and MASCOT-Skyline. Each subplot uses different settings for the simulations: low or high migration rates, where the mean migration rate was 5 resp. 25. 250 or 500 samples per state, or proportional.

**Figure S10:**
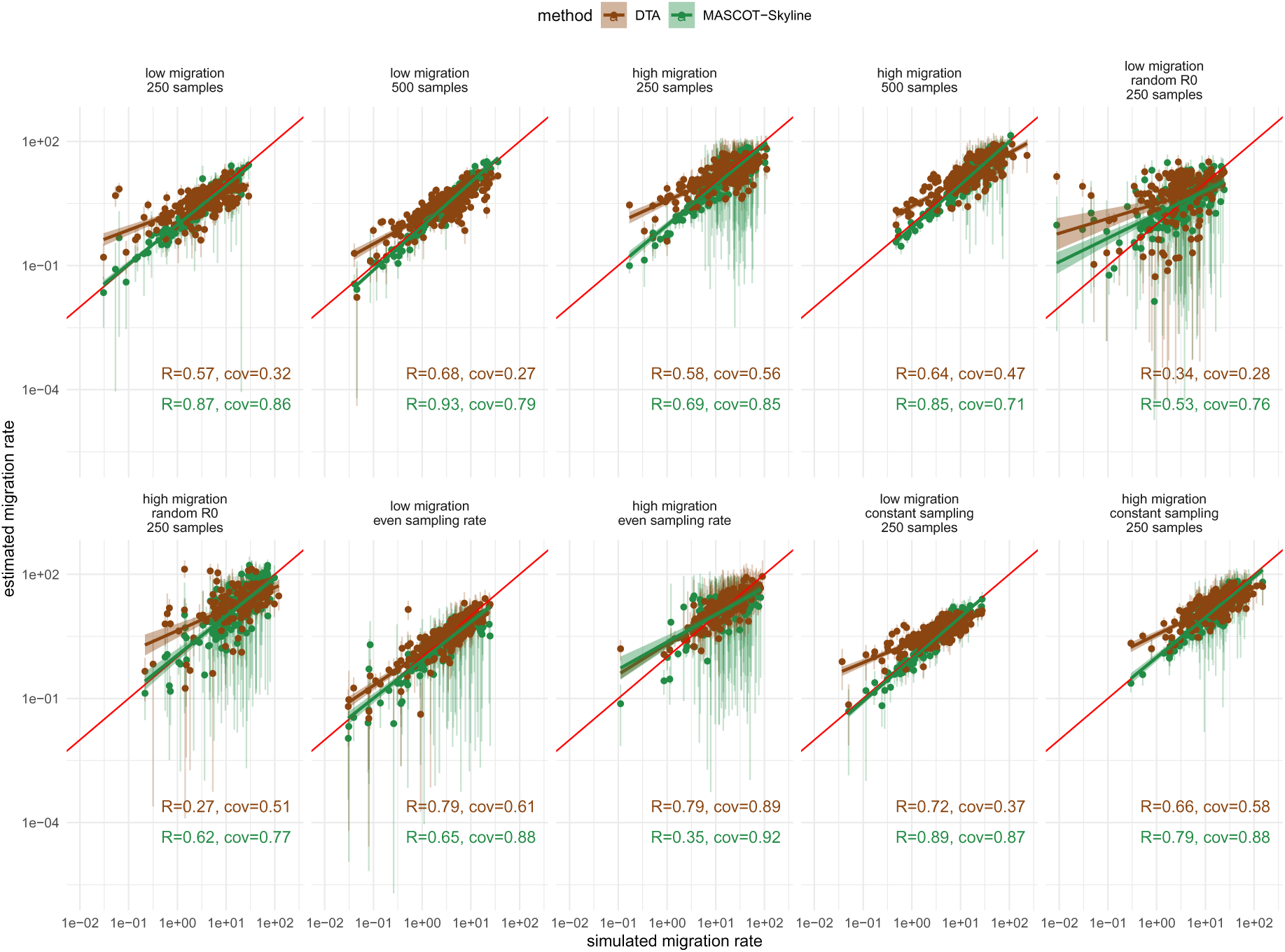
Correlations between the simulated and inferred migration rates for MASCOT-Skyline and DTA. Here, we show the simulated (x-axis) and estimated (y-axis) migration rates using simulations under a two-state SIR model. The dots show the median estimate, and the error bars show the 95% highest posterior density (HPD) interval. The person correlation coefficients (R) are calculated separately for MASCOT-Skyline and DTA. The coverage of the true value by the 95% HPD is shown after cov. The coefficients are calculated between the simulated values and the median estimates. Each subplot uses different settings for the simulations: low or high migration rates, where the mean migration rate was 5 resp. 25. 250 or 500 samples per state, or proportional and constant sampling.

**Figure S11:**
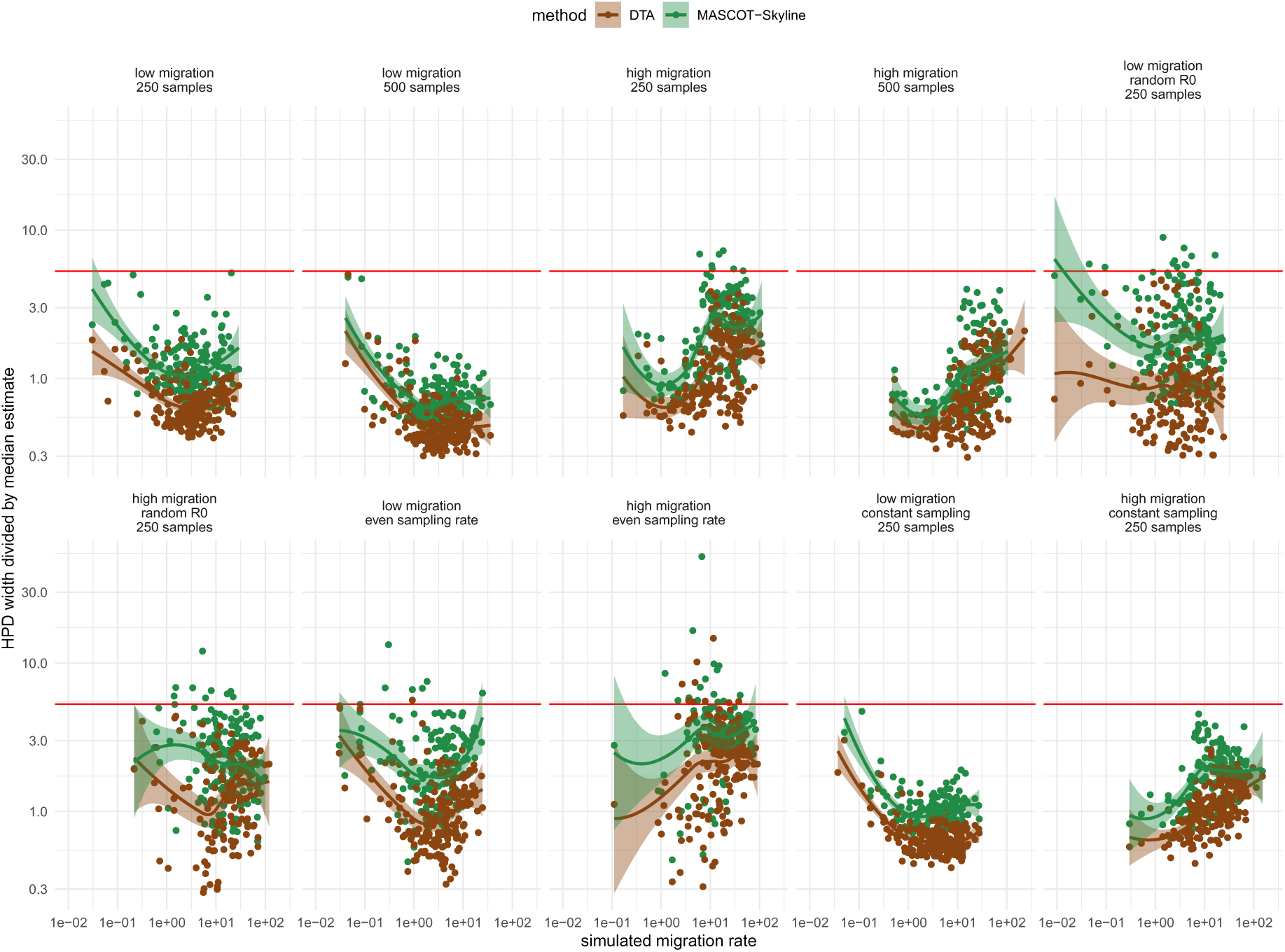
Relative HPD interval width for MASCOT-Skyline and DTA. Here, we show the simulated (x-axis) and estimated (y-axis) migration rates using simulations under a two-state SIR model. The dots show the difference between the upper and lower bound of the 95% highest posterior density interval divided by the median estimate. The red horizontal line shows the line for the upper and lower bound of the 95% interval of an exponential distribution used as a prior on the migration rates.

**Figure S12:**
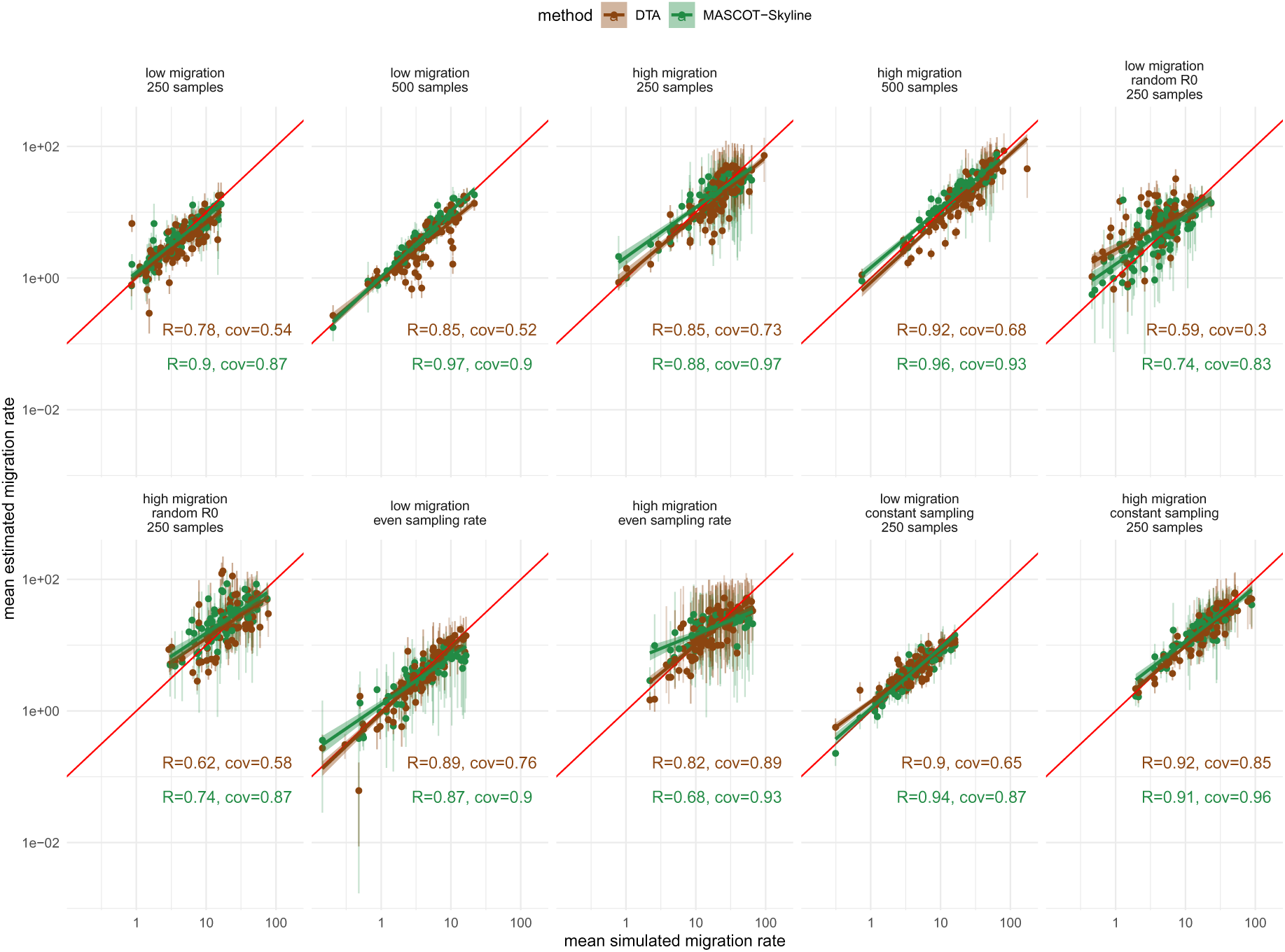
Estimation of the mean migration rate from two state SIR simulations. Here, we show the simulated (x-axis) and estimated (y-axis) migration rates using simulations under a two-state SIR model. The dots show the median estimate, and the error bars show the 95% highest posterior density (HPD) interval. The Pearson correlation coefficients (R) are calculated independently for MASCOT-Skyline and DTA, are shown in the top left corner of each plot, and are computed between the log of the true value and the log of the median estimate. We additionally show how often the 95% HPD interval covers the true value (cov). The coefficients are calculated between the simulated values and the median estimates. Each subplot uses different settings for the simulations, i.e., low or high migration rates, where the mean migration rate was 5 resp. 25. 250 or 500 samples per state, or proportional and constant sampling.

**Figure S13:**
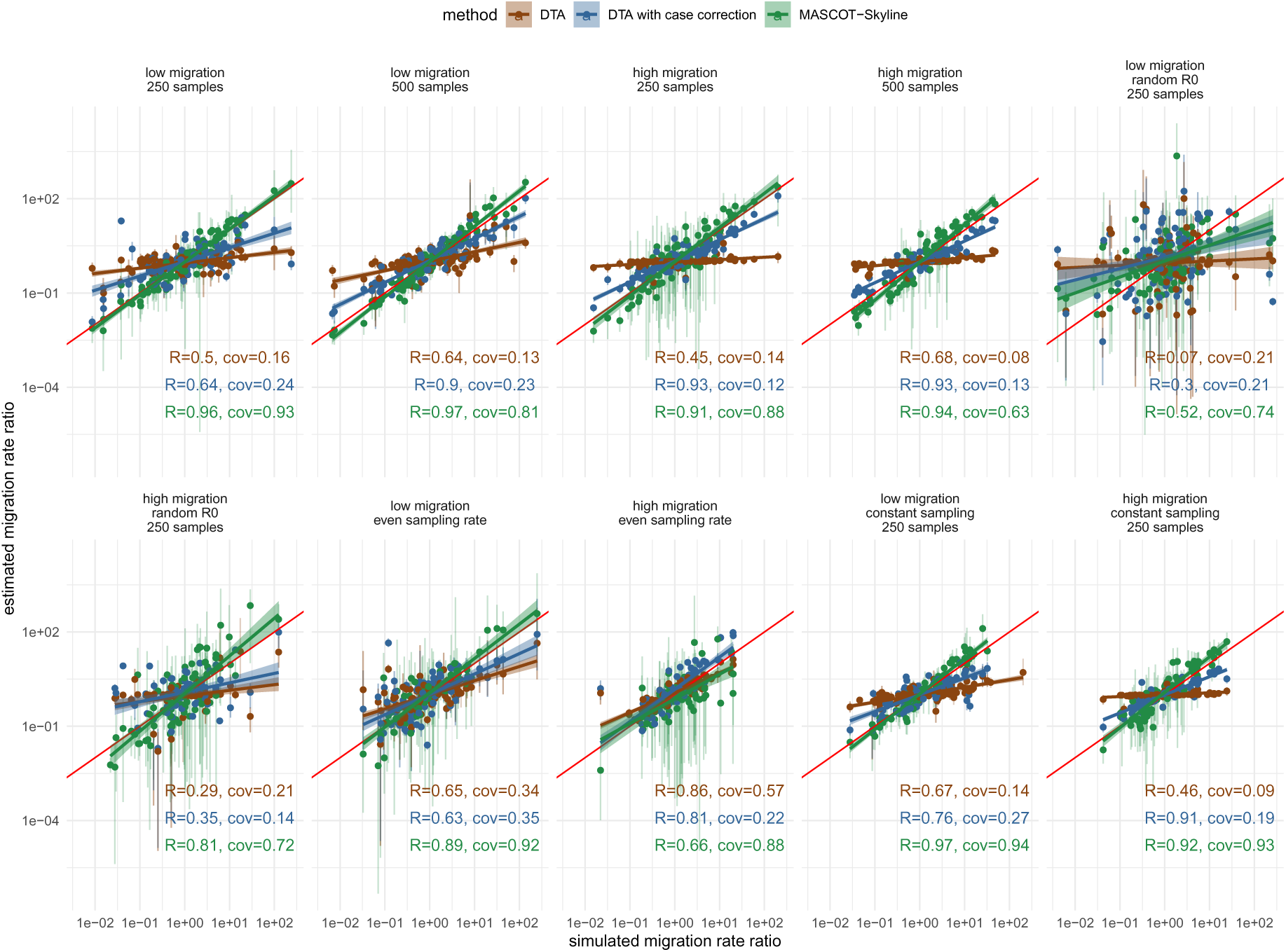
Simulated and inferred migration rate ratios using two state SIR simulations for MASCOT-Skyline and DTA. Here, we compare the simulated migration rate ratio to the estimated ratio of migration rates between the two states in the SIR model. The migration rate estimates are shown for DTA, MASCOT-Skyline, and DTA with case correction, where we multiply the ratio of migration rates with the ratio of cumulative incidence over the simulations to correct for differences in population size. The dots show the median estimate of the migration ratios, and the error bars show the 95% highest posterior density (HPD) interval. The Pearson correlation coefficients (R) are calculated independently for MASCOT-Skyline and DTA and DTA with case correction. The correlation coefficients are computed between the log of the true value and the log of the median estimate. We additionally show how often the 95% HPD interval covers the true value (cov). Each subplot uses different settings for the simulations, that is, low or high migration rates, where the mean migration rate was 5 resp. 25. 250 or 500 samples per state, proportional, and constant sampling.

**Figure S14:**
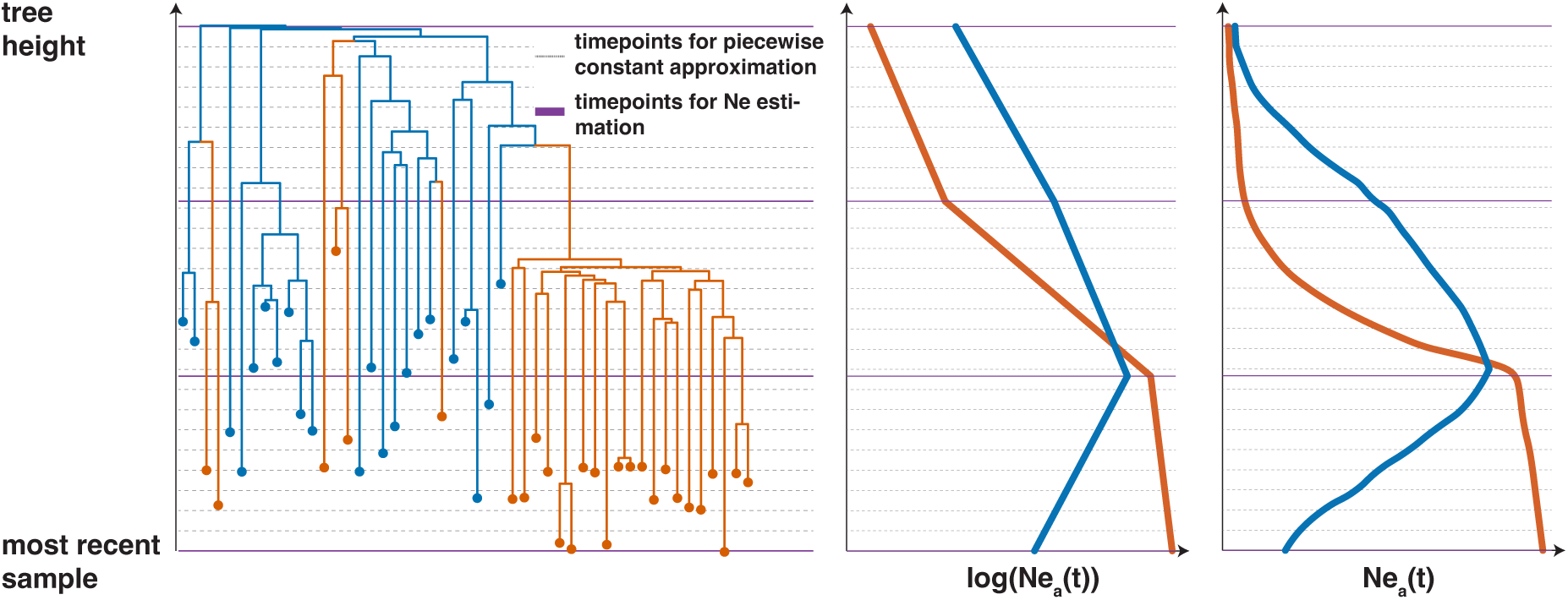
Description of how the effective population sizes are described over time. Each location in the dataset has its own population size trajectory. The population size trajectory is considered between the most recent sampled individual (mrsi) and the tree’s root. Here, we consider two more effective population sizes (Ne) between these two points in time. In this case, we estimate four Ne’s per location, with any number of Ne’s possible. Between the four points where we estimate the Ne, we assume that the Ne to change through exponential growth or decline. For the log of the Ne, that means we are using linear interpolation.

## References

Baele, G., Lemey, P., Rambaut, A., and Suchard, M. A. (2017). Adaptive MCMC in Bayesian phylogenetics: an application to analyzing partitioned data in BEAST. Bioinformatics, 33(12), 1798–1805.

Bahl, J., Nelson, M. I., Chan, K. H., Chen, R., Vijaykrishna, D., Halpin, R. A., Stockwell, T. B., Lin, X., Wentworth, D. E., Ghedin, E., et al. (2011). Temporally structured metapopulation dynamics and persistence of influenza a h3n2 virus in humans. Proceedings of the National Academy of Sciences, 108(48), 19359–19364.

Bedford, T., Cobey, S., Beerli, P., and Pascual, M. (2010). Global migration dynamics underlie evolution and persistence of human influenza a (h3n2). PLoS Pathog, 6(5), e1000918.

Bedford, T., Riley, S., Barr, I. G., Broor, S., Chadha, M., Cox, N. J., Daniels, R. S., Gunasekaran, C. P., Hurt, A. C., Kelso, A., et al. (2015). Global circulation patterns of seasonal influenza viruses vary with antigenic drift. Nature, 523(7559), 217–220.

Beerli, P. (2004). Effect of unsampled populations on the estimation of population sizes and migration rates between sampled populations. Molecular ecology, 13(4), 827–836.

Black, A., Moncla, L. H., Laiton-Donato, K., Potter, B., Pardo, L., Rico, A., Tovar, C., Rojas, D. P., Longini, I. M., Halloran, M. E., et al. (2019). Genomic epidemiology supports multiple introductions and cryptic transmission of zika virus in colombia. BMC infectious diseases, 19(1), 1–11.

Bouckaert, R., Vaughan, T. G., Barido-Sottani, J., Duchêne, S., Fourment, M., Gavryushkina, A., Heled, J., Jones, G., Kühnert, D., De Maio, N., et al. (2019). BEAST 2.5: An advanced software platform for Bayesian evolutionary analysis. PLoS computational biology, 15(4), e1006650.

Bouckaert, R. R. (2022). An efficient coalescent epoch model for Bayesian phylogenetic inference. Systematic Biology, 71(6), 1549–1560.

Brito, A. F., Semenova, E., Dudas, G., Hassler, G. W., Kalinich, C. C., Kraemer, M. U., Ho, J., Tegally, H., Githinji, G., Agoti, C. N., et al. (2022). Global disparities in sars-cov-2 genomic surveillance. Nature communications, 13(1), 7003.

De Maio, N., Wu, C.-H., O’Reilly, K. M., and Wilson, D. (2015). New routes to phylogeography: a Bayesian structured coalescent approximation. PLoS genetics, 11(8), e1005421.

De Maio, N., Wu, C.-H., and Wilson, D. J. (2016). Scotti: efficient reconstruction of transmission within outbreaks with the structured coalescent. PLoS computational biology, 12(9), e1005130.

Drummond, A. J. and Rambaut, A. (2007). BEAST: Bayesian evolutionary analysis by sampling trees. BMC evolutionary biology, 7(1), 1–8.

Drummond, A. J., Rambaut, A., Shapiro, B., and Pybus, O. G. (2005). Bayesian coalescent inference of past population dynamics from molecular sequences. Molecular biology and evolution, 22(5), 1185–1192.

Dudas, G., Carvalho, L. M., Rambaut, A., and Bedford, T. (2018). Mers-cov spillover at the camel-human interface. elife, 7, e31257.

Faria, N. R., Rambaut, A., Suchard, M. A., Baele, G., Bedford, T., Ward, M. J., Tatem, A. J., Sousa, J. D., Arinaminpathy, N., Pépin, J., et al. (2014). The early spread and epidemic ignition of hiv-1 in human populations. science, 346(6205), 56–61.

Faria, N. R., Quick, J., Claro, I., Theze, J., de Jesus, J. G., Giovanetti, M., Kraemer, M. U., Hill, S. C., Black, A., da Costa, A. C., et al. (2017). Establishment and cryptic transmission of zika virus in brazil and the americas. Nature, 546(7658), 406–410.

Faria, N. R., Kraemer, M. U., Hill, S. C., Góes de Jesus, J., Aguiar, R. d., Iani, F. C., Xavier, J., Quick, J., du Plessis, L., Dellicour, S., et al. (2018). Genomic and epidemiological monitoring of yellow fever virus transmission potential. Science, 361(6405), 894–899.

Gardy, J. L. and Loman, N. J. (2018). Towards a genomics-informed, real-time, global pathogen surveillance system. Nature Reviews Genetics, 19(1), 9–20.

Gibbons, C. L., Mangen, M.-J. J., Plass, D., Havelaar, A. H., Brooke, R. J., Kramarz, P., Peterson, K. L., Stuurman, A. L., Cassini, A., Fèvre, E. M., et al. (2014). Measuring underreporting and under-ascertainment in infectious disease datasets: a comparison of methods. BMC public health, 14(1), 1–17.

Gill, M. S., Lemey, P., Faria, N. R., Rambaut, A., Shapiro, B., and Suchard, M. A. (2013). Improving Bayesian population dynamics inference: a coalescent-based model for multiple loci. Molecular biology and evolution, 30(3), 713–724.

Grenfell, B. T., Pybus, O. G., Gog, J. R., Wood, J. L., Daly, J. M., Mumford, J. A., and Holmes, E. C. (2004). Unifying the epidemiological and evolutionary dynamics of pathogens. science, 303(5656), 327–332.

Grubaugh, N. D., Ladner, J. T., Kraemer, M. U., Dudas, G., Tan, A. L., Gangavarapu, K., Wiley, M. R., White, S., Thézé, J., Magnani, D. M., et al. (2017). Genomic epidemiology reveals multiple introductions of zika virus into the united states. Nature, 546(7658), 401–405.

Heller, R., Chikhi, L., and Siegismund, H. R. (2013). The confounding effect of population structure on Bayesian skyline plot inferences of demographic history. PloS one, 8(5), e62992.

Holmes, E. C. and Grenfell, B. T. (2009). Discovering the phylodynamics of rna viruses. PLoS computational biology, 5(10), e1000505.

Hudson, R. R. et al. (1990). Gene genealogies and the coalescent process. Oxford surveys in evolutionary biology, 7(1), 44.

Karcher, M. D., Palacios, J. A., Bedford, T., Suchard, M. A., and Minin, V. N. (2016). Quantifying and mitigating the effect of preferential sampling on phylodynamic inference. PLoS computational biology, 12(3), e1004789.

Kassambara, A. (2018). ggpubr:’ggplot2’based publication ready plots. R package version, page 2.

Kendall, D. G. (1948). On the generalized” birth-and-death” process. The annals of mathematical statistics, 19(1), 1–15.

Kingman, J. F. C. (1982). The coalescent. Stochastic processes and their applications, 13(3), 235–248.

Kühnert, D., Stadler, T., Vaughan, T. G., and Drummond, A. J. (2016). Phylodynamics with migration: a computational framework to quantify population structure from genomic data. Molecular biology and evolution, 33(8), 2102–2116.

Layan, M., Müller, N. F., Dellicour, S., De Maio, N., Bourhy, H., Cauchemez, S., and Baele, G. (2023). Impact and mitigation of sampling bias to determine viral spread: evaluating discrete phylogeography through ctmc modeling and structured coalescent model approximations. Virus Evolution, 9(1), vead010.

Lemey, P., Rambaut, A., Drummond, A. J., and Suchard, M. A. (2009). Bayesian phylogeography finds its roots. PLoS computational biology, 5(9), e1000520.

Lemey, P., Rambaut, A., Bedford, T., Faria, N., Bielejec, F., Baele, G., Russell, C. A., Smith, D. J., Pybus, O. G., Brockmann, D., et al. (2014). Unifying viral genetics and human transportation data to predict the global transmission dynamics of human influenza h3n2. PLoS pathogens, 10(2), e1003932.

Maddison, W. P., Midford, P. E., and Otto, S. P. (2007). Estimating a binary character’s effect on speciation and extinction. Systematic biology, 56(5), 701–710.

Merker, M., Blin, C., Mona, S., Duforet-Frebourg, N., Lecher, S., Willery, E., Blum, M. G., Rüsch-Gerdes, S., Mokrousov, I., Aleksic, E., et al. (2015). Evolutionary history and global spread of the mycobacterium tuberculosis beijing lineage. Nature genetics, 47(3), 242–249.

Minin, V. N., Bloomquist, E. W., and Suchard, M. A. (2008). Smooth skyride through a rough skyline: Bayesian coalescent-based inference of population dynamics. Molecular biology and evolution, 25(7), 1459–1471.

Müller, N. F., Rasmussen, D. A., and Stadler, T. (2017). The structured coalescent and its approximations. Molecular biology and evolution, 34(11), 2970–2981.

Müller, N. F., Rasmussen, D., and Stadler, T. (2018). Mascot: Parameter and state inference under the marginal structured coalescent approximation. Bioinformatics, 34(22), 3843–3848.

Müller, N. F., Dudas, G., and Stadler, T. (2019). Inferring time-dependent migration and coalescence patterns from genetic sequence and predictor data in structured populations. Virus evolution, 5(2), vez030.

Müller, N. F., Wagner, C., Frazar, C. D., Roychoudhury, P., Lee, J., Moncla, L. H., Pelle, B., Richardson, M., Ryke, E., Xie, H., et al. (2021). Viral genomes reveal patterns of the sars-cov-2 outbreak in washington state. Science Translational Medicine, 13(595), eabf0202.

Notohara, M. (1990). The coalescent and the genealogical process in geographically structured population. Journal of mathematical biology, 29, 59–75.

Plummer, M., Best, N., Cowles, K., Vines, K., et al. (2006). Coda: convergence diagnosis and output analysis for mcmc. R news, 6(1), 7–11.

Popinga, A., Vaughan, T., Stadler, T., and Drummond, A. J. (2015). Inferring epidemiological dynamics with Bayesian coalescent inference: the merits of deterministic and stochastic models. Genetics, 199(2), 595–607.

Pybus, O., Drummond, A., Nakano, T., Robertson, B., and Rambaut, A. (2003). The epidemiology and iatrogenic transmission of hepatitis C virus in Egypt: a Bayesian coalescent approach. Molecular biology and evolution, 20(3), 381–387.

Rambaut, A., Pybus, O. G., Nelson, M. I., Viboud, C., Taubenberger, J. K., and Holmes, E. C. (2008). The genomic and epidemiological dynamics of human influenza a virus. Nature, 453(7195), 615–619.

Slatkin, M. (2005). Seeing ghosts: the effect of unsampled populations on migration rates estimated for sampled populations. Molecular ecology, 14(1), 67–73.

Stadler, T. and Bonhoeffer, S. (2013). Uncovering epidemiological dynamics in heterogeneous host populations using phylogenetic methods. Philosophical Transactions of the Royal Society B: Biological Sciences, 368(1614), 20120198.

Stadler, T., Kühnert, D., Bonhoeffer, S., and Drummond, A. J. (2013). Birth–death skyline plot reveals temporal changes of epidemic spread in hiv and hepatitis c virus (hcv). Proceedings of the National Academy of Sciences, 110(1), 228–233.

Stolz, U., Stadler, T., Müller, N. F., and Vaughan, T. G. (2022). Joint inference of migration and reassortment patterns for viruses with segmented genomes. Molecular biology and evolution, 39(1), msab342.

Strimmer, K. and Pybus, O. G. (2001). Exploring the demographic history of dna sequences using the generalized skyline plot. Molecular Biology and Evolution, 18(12), 2298–2305.

Suchard, M. A., Lemey, P., Baele, G., Ayres, D. L., Drummond, A. J., and Rambaut, A. (2018). Bayesian phylogenetic and phylodynamic data integration using beast 1.10. Virus evolution, 4(1), vey016.

Takahata, N. (1988). The coalescent in two partially isolated diffusion populations. Genetics Research, 52(3), 213–222.

Vaughan, T. G. and Drummond, A. J. (2013). A stochastic simulator of birth–death master equations with application to phylodynamics. Molecular biology and evolution, 30(6), 1480–1493.

Volz, E. M. (2012). Complex population dynamics and the coalescent under neutrality. Genetics, 190(1), 187–201.

Volz, E. M. and Didelot, X. (2018). Modeling the growth and decline of pathogen effective population size provides insight into epidemic dynamics and drivers of antimicrobial resistance. Systematic Biology, 67(4), 719–728.

Volz, E. M. and Siveroni, I. (2018). Bayesian phylodynamic inference with complex models. PLoS computational biology, 14(11), e1006546.

Volz, E. M., Kosakovsky Pond, S. L., Ward, M. J., Leigh Brown, A. J., and Frost, S. D. (2009). Phylodynamics of infectious disease epidemics. Genetics, 183(4), 1421–1430.

Volz, E. M., Koelle, K., and Bedford, T. (2013). Viral phylodynamics. PLoS computational biology, 9(3), e1002947.

Vos, T., Lim, S. S., Abbafati, C., Abbas, K. M., Abbasi, M., Abbasifard, M., Abbasi-Kangevari, M., Abbastabar, H., Abd-Allah, F., Abdelalim, A., et al. (2020). Global burden of 369 diseases and injuries in 204 countries and territories, 1990–2019: a systematic analysis for the global burden of disease study 2019. The Lancet, 396(10258), 1204–1222.

Wickham, H. (2016). ggplot2: elegant graphics for data analysis. Springer.

Worobey, M., Watts, T. D., McKay, R. A., Suchard, M. A., Granade, T., Teuwen, D. E., Koblin, B. A., Heneine, W., Lemey, P., and Jaffe, H. W. (2016). 1970s and ‘patient 0’hiv-1 genomes illuminate early hiv/aids history in north america. Nature, 539(7627), 98–101.

Yu, G., Smith, D. K., Zhu, H., Guan, Y., and Lam, T. T.-Y. (2017). ggtree: an r package for visualization and annotation of phylogenetic trees with their covariates and other associated data. Methods in Ecology and Evolution, 8(1), 28–36.

